# High-throughput variant libraries and machine learning enable robust target-site selection for toehold-switch design in Detectrons

**DOI:** 10.64898/2026.10.09.758053

**Authors:** Jihoon Han, Seth L. Shipman

## Abstract

Programmable RNA sensors enable sequence-specific detection of RNA inputs for diagnostics, genetic circuit control, and multiplexed molecular readouts. Although readily programmable, their performance varies substantially with the target site selected within a transcript, making target-site selection a major challenge. Here, we report DeSTAR (Detectron Sensor TARget-site ranker), a machine-learning-assisted target-site ranking model for programmable RNA sensors. To train this model, we generated transcript-wide, single-nucleotide-resolution tiling sensor libraries across six diverse RNA targets, enabling generalization to be tested across transcripts rather than within a single RNA. In leave-one-transcript-out evaluation, DeSTAR’s top-ranked sites were enriched for high-performing sensors in held-out transcripts, driven predominantly by interpretable sequence grammar in the sensor stem rather than transcript-specific patterns. Prospectively, on an unseen SARS-CoV-2 transcript, the model’s top-ranked sites yielded working sensors roughly five-fold more often than random selection. These results support DeSTAR as a generalizable strategy for target-site selection in RNA sensors across transcripts.

## INTRODUCTION

Programmable RNA sensors such as toehold switches can be rationally designed to detect virtually any RNA sequence, and this programmability has enabled a new generation of low-cost, field-deployable diagnostics^1–8^. In principle, a sensor can be designed against any region of a target transcript. In practice, however, performance varies dramatically depending on which site is chosen, and most candidate sites yield poorly performing sensors — making target-site selection a central bottleneck in RNA sensor engineering, analogous to the guide-RNA selection problem that machine learning has helped solve in CRISPR systems^9^.

The original toehold switches already revealed some determinants of this variation: Green et al. found that performance depends on the base-pairing composition of the switch stem — favoring weak (A·U) pairs at the loop-proximal top and two strong (G·C) pairs out of three at the base — though only as a count-based rule from 168 designed switches^1^. To make such rules predictive at scale, subsequent efforts turned to machine learning trained on large experimental datasets. Two companion studies characterized a shared library of 91,534 toehold switches (from 244,000 designed sequences) by high-throughput flow-seq and showed that deep neural networks trained on switch sequence predict function more accurately than thermodynamic or kinetic models^10,11^. These studies established sequence-to-function learning as a powerful paradigm for riboregulatory design. Their models, however, operate on the switch sequence itself. Because each candidate was characterized against a short cognate trigger rather than a target site within a full-length transcript, the structural context and accessibility of the target within its parent RNA — which depend on transcript-wide folding — were absent from these datasets and could not be modeled. These datasets are also based entirely on fluorescent protein output — the only toehold-switch readout available at the time — so whether such rules extend to sensors with fundamentally different output modalities had not been addressed.

More recently, Robson and Green reframed the problem as one of target-aware design: rather than scoring the switch alone, they modeled the folding and accessibility of the target site within its transcript, using a switch backbone that minimizes stem variation to isolate target-side effects^12^. By pairing a tiled toehold-switch library with biophysical features of both sensor and target — local base-pairing probabilities, flanking-window free energies, and even codon-usage effects on co-transcriptional folding — their Toehold-VISTA (versatile in-silico RNA-targeting analysis) framework predicted sensor performance more accurately than existing design tools and, in a prospective test, selected high-performing sensors for an unseen SARS-CoV-2 target. In their model, sensor thermodynamic stability — the free energy of the ON and OFF states — remained the dominant determinant, with target-site accessibility emerging as a key additional determinant. The study’s scope, however, was bounded by the experimental design: a single transcript (mCherry) tiled at three-nucleotide intervals, yielding roughly 200 sensors, each measured in a separate flow-cytometry sample. Whether rules learned from this single transcript generalize across structurally and biologically diverse RNAs therefore remains to be established. Moreover, the non-pooled readout limits the throughput for surveying many transcripts at single-nucleotide resolution.

Here, we report DeSTAR (Detectron Sensor TARget-site ranker), a machine-learning-assisted target-site ranking model that addresses these limitations directly. DeSTAR is built on the Detectron, a DNA-barcode-output sensor architecture we recently developed^13^. In a Detectron, a toehold switch is fused to a retron so that, in the absence of the target RNA, the switch hairpin sequesters the retron’s a2 region, disrupting the a1/a2 complementarity required to prime reverse transcription and blocking DNA synthesis; binding of the target RNA drives strand displacement that restores a1/a2 pairing and triggers synthesis of a sequenceable DNA barcode (**Fig. 1a**). Because the output is a DNA barcode rather than a protein signal, many sensors can be pooled and quantified together in a single sequencing measurement. This pooled readout made it feasible to tile candidate sites at single-nucleotide resolution across six transcripts spanning three biological classes: two bacteriophage transcripts (T5 gp8, T7 gp10A), two human-pathogenic viral transcripts — NS4A-NS4B region of Zika virus (ZIKV) and the envelope (E) region of dengue virus serotype 1 (DENV-1) — and two human transcripts (*PGK1* and *ENO1*), yielding 7,383 sensors in total (**Fig. 1b**). These transcript-wide libraries revealed broad site-to-site variation in sensor performance, driven predominantly by a positional base-pairing grammar at the base of the switch stem. By integrating these features, we trained a model that ranked candidate target sites within individual RNAs (**Fig. 1c**) and, in a leave-one-transcript-out evaluation, generalized to all held-out transcripts, enriching for high-performing sites well above chance. Finally, in a prospective test on an unseen SARS-CoV-2 transcript, the model’s top-ranked sites yielded working sensors far more often than randomly chosen sites. Together, these results establish a generalizable framework for selecting target sites for programmable RNA sensors across diverse transcripts.

**Figure 1.**
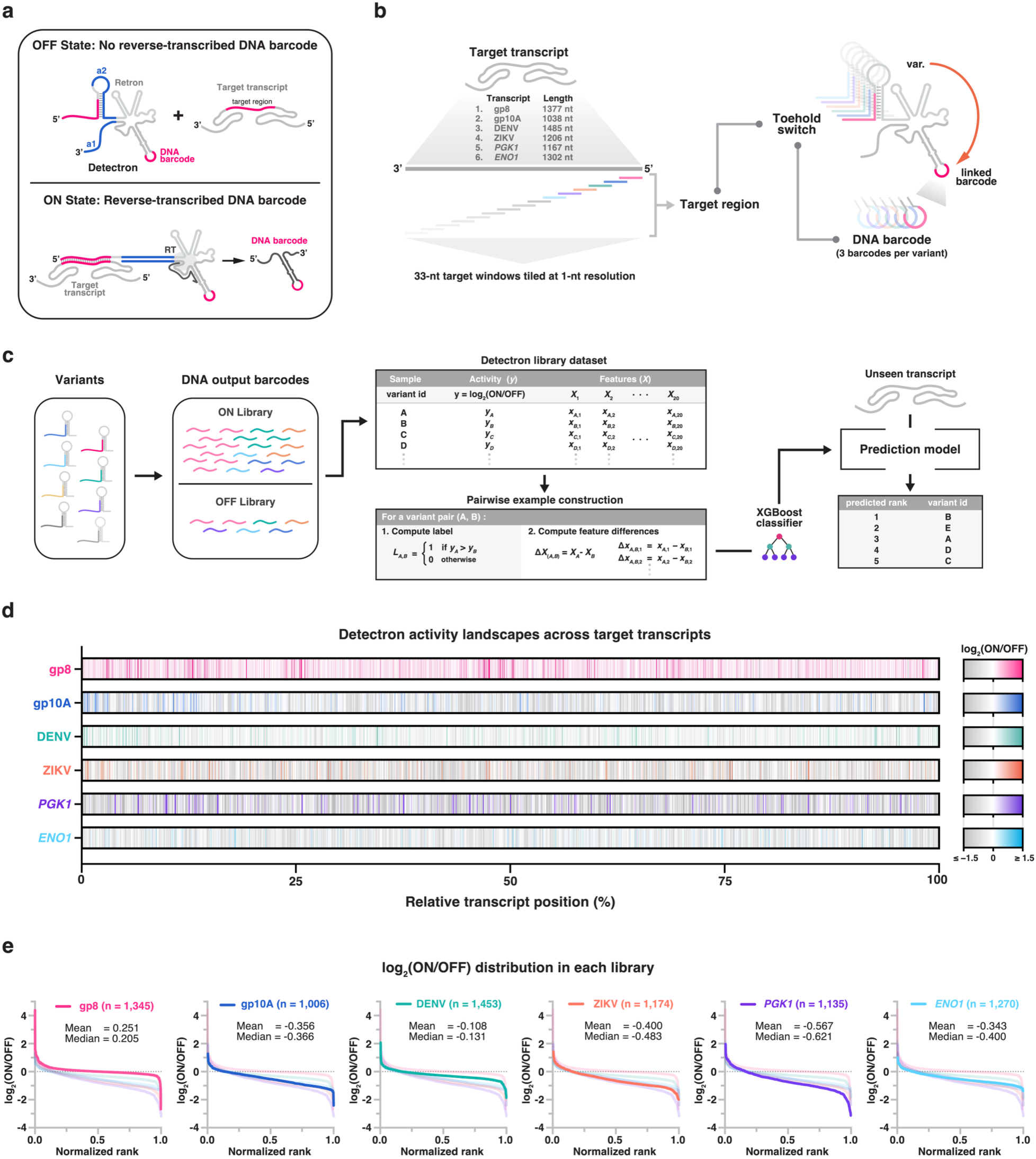
Transcript-wide Detectron tiling libraries reveal target-site-dependent variation in RNA sensor activity. **a**, Detectron sensing mechanism. In the OFF state (no target transcript), no DNA barcode is produced. In the ON state, base-pairing of the transcript’s target region opens the toehold switch, enabling the retron reverse transcriptase (RT) to synthesize a reverse-transcribed DNA barcode. **b**, Library design. Each of six target transcripts (identities and lengths listed) was tiled into overlapping 33-nt target windows at single-nucleotide resolution. Each window defines one toehold-switch variant carrying a linked DNA barcode (three barcodes per variant), yielding a pooled sensor library spanning the full transcript. **c**, Dataset and ranking model. ON- and OFF-state barcodes were sequenced and their abundances quantified; sensor activity is quantified as y = log_2_(ON/OFF), where ON/OFF is the ratio of barcode abundance in the ON versus OFF states. Each variant is described by 20 sequence/structure features (*X*). Variant pairs (A,B) were labelled by which had higher activity (*L*_A,B_ = 1 if y_A_ > y_B_, else 0) and encoded as feature differences (Δ*X* = *X*_A_ – *X*_B_); a pairwise XGBoost classifier trained on these differences ranks candidate target sites in an unseen transcript. **d**, Detectron activity landscapes. log_2_(ON/OFF) of every tiled target site, arranged by relative position along each transcript (0-100%). Color indicates log_2_(ON/OFF), clipped at ± 1.5 (values below −1.5 or above +1.5 are shown at the scale limits). **e**, Distribution of log_2_(ON/OFF) within each library: each variant’s activity plotted against its normalized rank (variants sorted from highest to lowest activity; rank scaled 0-1). Mean and median are indicated. Faint background curves show the other five libraries for comparison. *n* = number of tiled target sites per transcript.

## RESULTS

### Transcript-wide Detectron tiling libraries reveal target-site-dependent variation in RNA sensor activity

To map how sensor activity varies with target-site position within a transcript, we built transcript-wide tiling libraries of Detectron sensors (**Fig. 1a**) across six diverse RNA targets. The six transcripts span three biological classes and differ in length, GC content, and predicted folding stability (**Fig. 1b, Supplementary Table 1**). Each transcript was tiled at single-nucleotide resolution with a 33-nt window, so that every 33-nt region became the target of one Detectron sensor, yielding 1,006-1,453 target sites per transcript and 7,383 sensors in total (**Fig. 1b**). For each target site, we designed a Detectron in which a unique DNA barcode, encoded in the retron msd (multicopy single-stranded DNA–encoding region), identifies that site (**Fig. 1b**). To control for sequence-specific effects of individual barcodes on reverse-transcribed DNA (RT-DNA) production, each sensor was represented by three distinct barcodes (**Fig. 1b**), giving 22,149 barcoded constructs.

Each library was assayed as paired ON and OFF sublibraries, in which the cognate target transcript was constitutively expressed or absent, respectively (**Fig. 1c**). For each sensor, we sequenced its barcoded RT-DNA and determined the relative barcode abundance in the ON and OFF states, taking the ON/OFF ratio — the fold-change in DNA-barcode output upon target recognition — as the measure of sensor performance, reported as log_2_(ON/OFF) (hereafter “activity”)^13^. The ON/OFF ratio is the figure of merit for a sensor because it captures the separation between the target-present and target-absent states: a useful sensor must produce substantially more output when its target is present than when it is absent, and it is this ratio — not the absolute ON-state output — that determines whether target presence can be reliably distinguished from background. Using this metric, we found that within every transcript, activity varied markedly from site to site, with a minority of positions yielding high-performing sensors interspersed among many low-activity sites and no consistent positional pattern (**Fig. 1d**). Remarkably, this variation occurred even between immediately adjacent target sites: shifting a target site by a single nucleotide could turn a high-performing sensor into an inactive one (**Extended Data Fig. 1a-f**). Indeed, a single-nucleotide shift changed activity almost as much as moving to a randomly chosen site elsewhere in the transcript (**Extended Data Fig. 1g**). These single-nucleotide differences exceeded replicate measurement noise in five of six transcripts — all except DENV, where they did not (sign test, n.s.) — though their magnitude varied across libraries.

Because high-performing sites can lie immediately beside inactive ones, tiled testing of target sites at single-nucleotide resolution was essential: coarser spacing would have skipped many of the rare functional sites, and the best target site for a transcript could have been missed entirely. The magnitude and spread of activity also differed substantially between transcripts (**Fig. 1e**). gp8 was the only transcript centered above ON/OFF = 1 (median log_2_ = 0.205) and reached the highest peak activities (up to log_2_ = 4.6), yet its bulk distribution was the most tightly clustered. The other transcripts were centered below 1 and varied in spread, with *PGK1* both the lowest (median log_2_ = −0.62) and the most dispersed — indicating that both target-site choice and transcript identity shape sensor performance.

### A bottom-stem base-pairing grammar distinguishes high-from low-performing sensors

Because activity varied from site to site with no consistent positional pattern, we asked whether sequence features shared across transcripts distinguish high-from low-performing target sites. The Detectron switch presents a 33-nt variable region — the reverse complement of the target site — comprising a 25-nt single-stranded toehold that binds the target and an 8-nt segment that fold into the switch stem, which is closed by three invariant weak base pairs (**Fig. 2a**). Within each transcript we ranked sites by measured activity, pooled the top and bottom 15% (high- and low- performing, respectively) across all six transcripts (*n* = 1,107 per group), and compared their base composition position by position as an enrichment logo [log2(High/Low)] (**Fig. 2b**). Sequence enrichment was sharply localized to the switch stem: the single-stranded toehold (positions 1-25) was essentially unbiased, whereas the eight base pairs from the base of the stem (positions 26-33) discriminated strongly between high- and low-performing sensors (**Fig. 2c**).

**Figure 2.**
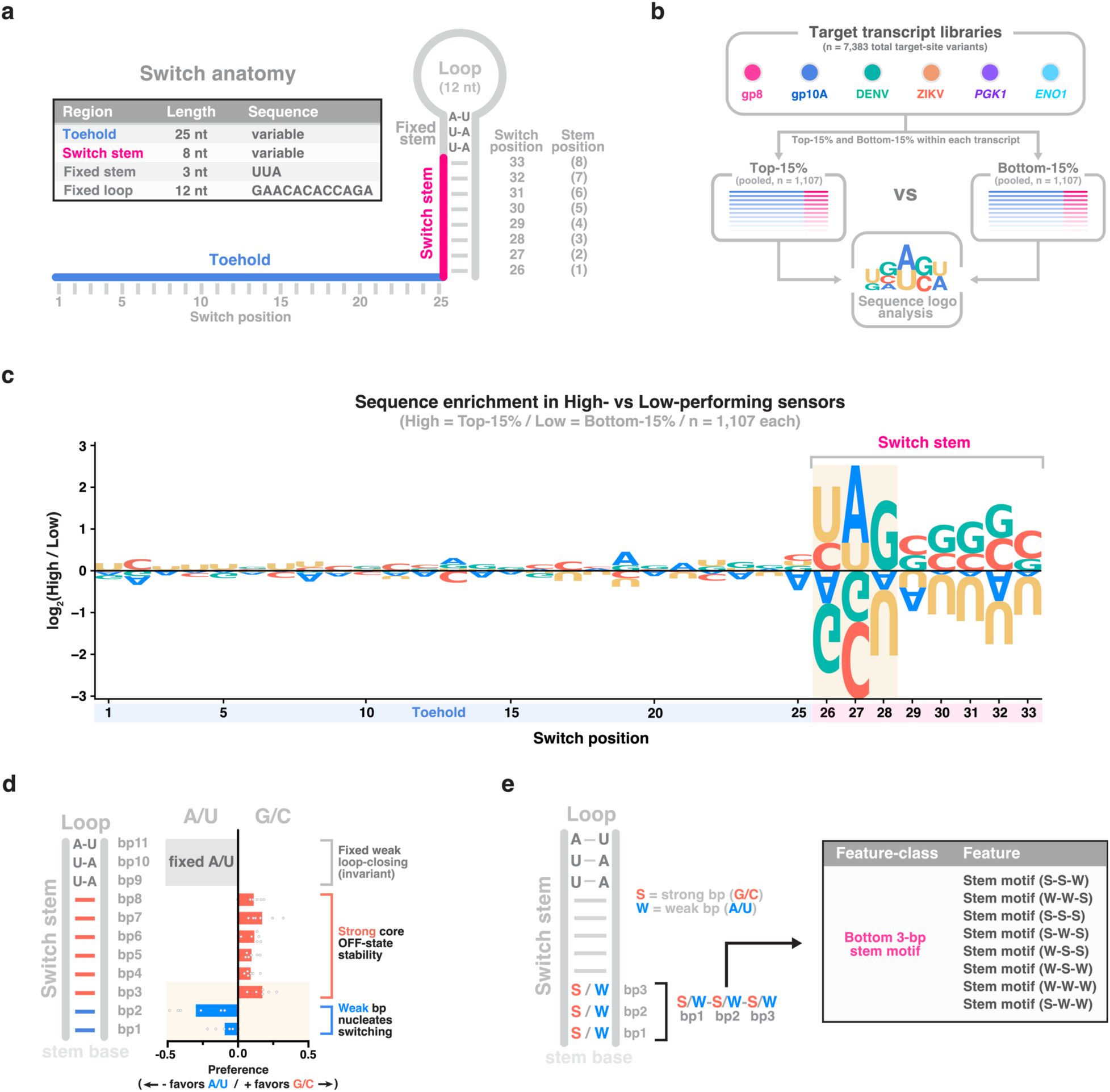
A bottom-stem base-pairing grammar distinguishes high- from low-performing sensors. **a**, Anatomy of the Detectron switch. Each sensor presents a 33-nt variable region — a 25-nt single-stranded toehold that binds the target and an 8-bp switch stem (switch positions 26-33, bp1 [toehold-proximal] to bp8 [loop-proximal]) — by three invariant weak (A-U) loop-closing base pairs. The variable sequence is the reverse complement of the target site. **b**, Sequence-logo analysis workflow. Across six transcript libraries (gp8, gp10A, DENV, ZIKV, *PGK1*, *ENO1*; *n* = 7,383 target-site variants in total), sensors were ranked by measured activity within each transcript, and each transcript’s top and bottom 15% were pooled across all six (*n* = 1,107 per group) for sequence-logo comparison. **c**, Sequence-enrichment logo comparing high- (top 15%) and low-performing (bottom 15%) sensors. Letter height is the log_2_ ratio of base frequency [log_2_(High/Low)]; letters above the axis are enriched in high performers, below in low. Enrichment is confined to the switch stem (positions 25-33); the toehold (1-25) is essentially unbiased. bp1-3 (positions 26-28) shaded; *n* = 1,107 per group. **d**, Position-resolved base-pairing preference across the switch stem. Preference, Δmean log_2_(ON/OFF) between variants with a strong versus weak pair at each position (positive favors G·C). Variable positions bp1-bp8 show a weak-base/strong-core polarity while bp9-bp11 are invariant A·U pairs (grey). *n* = 6 transcripts. Bars, mean across six transcripts. **e**, Definition of the bottom-stem motif feature class used in the ranking model. The strong (S = G/C) or weak (W = A/U) identity of the three toehold-proximal base pairs (bp1-bp3) defines 2^3^ = 8 motif classes, encoded as eight features.

The discriminating signal followed a polar architecture along the stem. At the level of pairing strength (A/U vs G/C), the two base pairs at the stem base (bp1-2, positions 26-27) leaned toward weak A/U pairs in high-performing sensors, whereas the six pairs above them (bp3-8, positions 28-33) favored strong G/C pairs (**Fig. 2c,d**); this weak-base/strong-core pattern was consistent across all six transcripts (**Extended Data Fig. 2**) and robust to the activity cut-off used to define high and low performers (**Extended Data Fig. 3**).

Within this weak-pair region, bp1 and bp2 carried different kinds of signal. Bp2 showed the strongest preference of any stem position and followed pairing strength: high-performing sensors favored a weak A/U pair, with A the most enriched (**Fig. 2c,d**). The terminal pair bp1, by contrast, showed little strength preference but a distinct base-identity signal, favoring pyrimidines — most strongly U, with even the strongly pairing C enriched over the weakly pairing A (**Fig. 2c**). This difference in kind most likely reflects the two positions’ distinct structural roles: bp2 remains paired and is read out through its pairing strength, whereas bp1, as the terminal pair at the single-stranded toehold junction, is prone to fraying^14^ and is therefore governed by base identity rather than by pairing strength.

Given this reproducible weak-base/strong-core architecture — the finer base identities above being mechanistic signatures rather than ranking determinants, as we confirm later — we encoded this region compactly as a bottom-stem motif feature class: the strong/weak identity of the first three base pairs (2^3^ = 8 features), included as one of five feature classes in the ranking model.

### A pairwise ranking model enriches for high-performing target sites across held-out transcripts

The bottom-stem grammar identifies one strong determinant of performance, but ranking target sites in a new transcript requires combining it with the remaining sequence and context features and testing whether such a model generalizes beyond the transcripts used to build it. We therefore trained a pairwise machine-learning model and assessed its generalization by leave-one-transcript-out (LOTO) cross validation (**Fig. 3a**). In each fold, the model was trained on five transcript libraries and used to rank the target sites of the held-out sixth; this was repeated for all six transcripts and averaged over ten random seeds. We represented each candidate target site by 20 features capturing sensor-intrinsic properties (bottom-stem sequence motif, hairpin and toehold folding energies) and transcript-specific context (target-site accessibility, GC content, and relative transcript position) (**Methods**; **Supplementary Table 2**). All features and per-replicate ON/OFF values for every sensor are provided in Supplementary Table 3. Because absolute ON/OFF ranges differed markedly between transcripts (**Fig. 1e**), we did not regress absolute activity; instead we framed site selection as a within-transcript ranking problem, which asks only which of two sites in the same transcript performs better and is therefore insensitive to transcript-level offsets (**Fig. 3a**). Specifically, for every pair of sites (A, B) within a transcript, we defined a binary label indicating whether site A outperformed site B: *L*_A,B_ = 1 when site A had higher measured activity than B (y_A_ > y_B_, where y = log_2_(ON/OFF)) and 0 otherwise. Each pair was represented by the difference of the two sites’ feature vectors, Δ*X*_(A,B)_ = *X*_A_ – *X*_B_, so that the model learned from relative rather than absolute feature values. A gradient-boosted decision-tree classifier^15^ (XGBoost) was trained on these pairwise examples to estimate P(A > B), the probability that site A outperforms site B — the outcome the label encodes (**Fig. 3a**). To rank the sites of the held-out transcript, each candidate was scored by its average predicted win probability against all other candidates in that transcript, and sites were ordered by this score.

**Figure 3.**
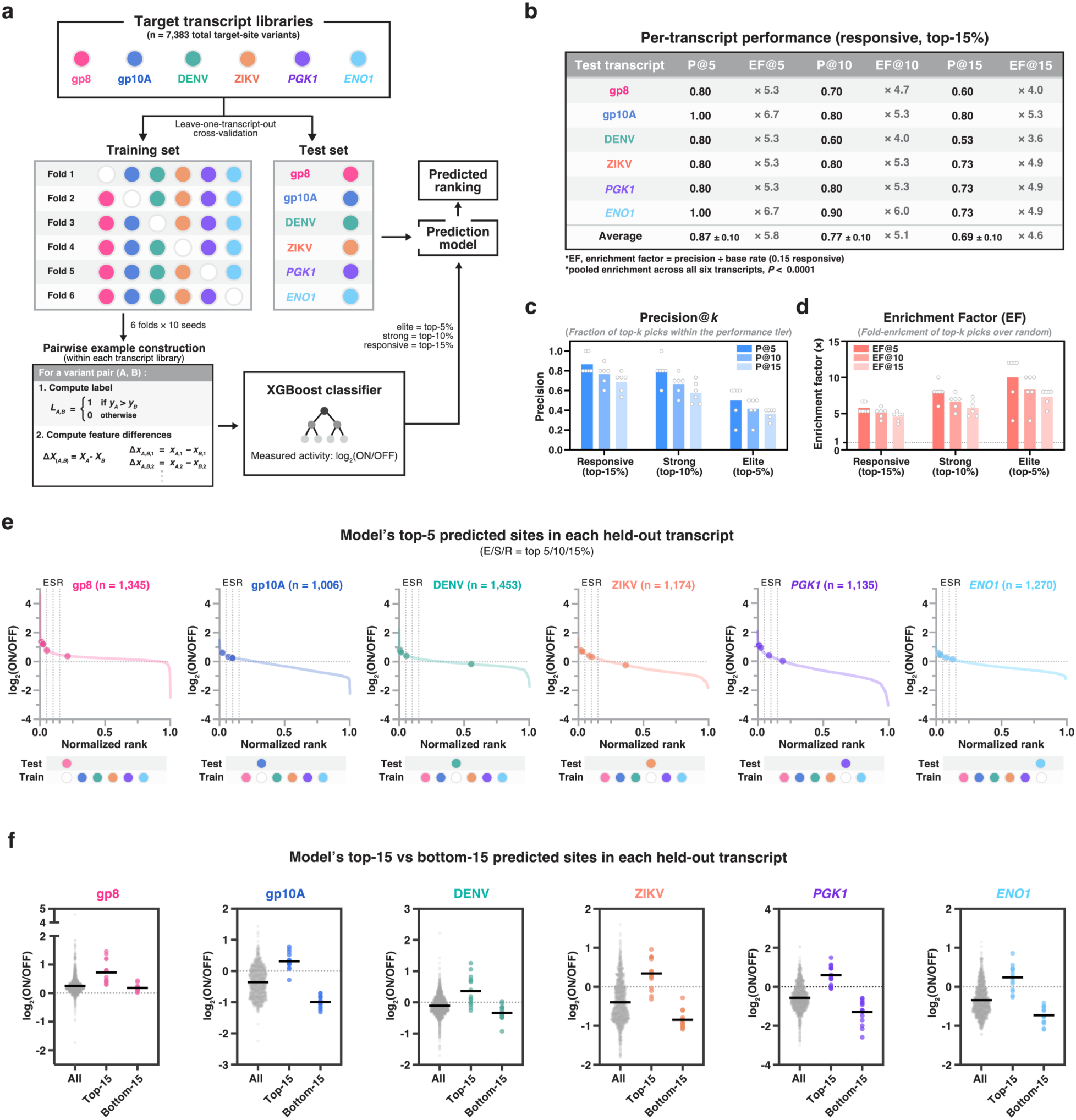
A pairwise ranking model enriches for high-performing target sites across held-out transcripts. **a**, Model overview. Six transcript-wide Detectron tiling libraries (gp8, gp10A, DENV, ZIKV, *PGK1*, *ENO1*; *n* = 7,383 total target-site variants) were evaluated by leave-one-transcript-out cross-validation: the model was trained on five libraries and tested on the held-out sixth, across all six held-out choices (6 folds × 10 random seeds). Within each training library, variant pairs (A, B) were labelled by which had higher measured activity (*L*_A,B_ = 1 if y_A_ > y_B_, else 0) and encoded as feature differences (Δ*X* = *X*_A_ – *X*_B_); an XGBoost classifier trained on these differences scored each candidate site in the held-out transcript by its mean pairwise-win probability against all other sites, and scores were averaged across the 10 seeds to give one consensus ranking per transcript. Activity was measured as log_2_(ON/OFF); performance tiers were defined per transcript by measured activity — responsive (top 15%), strong (top 10%), elite (top 5%). **b**, Per-transcript ranking performance at the responsive (top 15%) tier. Precision@*k* (P@*k*, fraction of the top-*k* predicted sites falling within the tier) and enrichment factor (EF = precision ÷ base rate, the fold-enrichment over random site selection) for *k* = 5, 10, 15. Each row reports one transcript (precision and EF computed from that transcript’s consensus ranking); the Average row is the mean ± s.d. across the six transcripts. Pooled enrichment across all six transcripts, *P* < 0.0001 (all *k*; hypergeometric enrichment test versus random site selection). **c**, Precision@*k* of the model’s top-*k* predicted sites at three performance tiers — responsive (top 15%), strong (top 10%) and elite (top 5%) — for *k* = 5, 10, and 15 (bars within each tier, dark to light). Bars, mean across six held-out transcripts. **d**, Enrichment factor (EF = precision ÷ tier base rate) for the same tiers and *k*, i.e. the fold-enrichment of the top-*k* predictions over random site selection (dashed line, EF = 1). Bars, mean across six held-out transcripts. Enrichment was significant at every tier and *k* (*P* < 0.0001; pooled hypergeometric enrichment test versus random site selection). **e**, Measured activity (log_2_(ON/OFF)) of every candidate site in each held-out transcript, ordered by measured activity (normalized rank), with the model’s five top-ranked predicted sites overlaid (filled circles). Dotted lines mark the elite, strong, and responsive tier boundaries (E/S/R = top 5%/10%/15%). n per transcript as indicated. **f**, Measured activity of all candidate sites (All), the model’s 15 top-ranked predicted sites (Top 15) and 15 bottom-ranked predicted sites (Bottom 15) in each held-out transcript. In all six transcripts, top-ranked sites had significantly higher activity than both the bottom-ranked sites and the all-site mean. (Mann-Whitney *U*-test, two-tailed; per-transcript statistics in Supplementary Table 4).

We quantified performance with precision@*k* — the fraction of the top-*k* predicted sites that fell within a defined performance tier — and the enrichment factor (EF = precision ÷ base rate), the fold-enrichment over random site selection. We assessed three tiers, defined per transcript by measured activity: responsive (top 15%), strong (top 10%), and elite (top 5%). Across all six held-out transcripts, the model’s top-ranked sites were strongly enriched for high-performing sensors. At the responsive tier, the predicted top 5 averaged 87% precision (EF = 5.8×), with enrichment sustained at deeper cut-offs (top 10: 77%, EF = 5.1×; top 15: 69%, EF = 4.6×) (**Fig. 3b**). This enrichment extended to the strictest tier: the predicted top-5 reached 10× enrichment for elite (top-5%) sites over random site selection (pooled hypergeometric *P* < 0.0001; **Supplementary Table 4**), so the model’s prioritization is not limited to the broader top-15% but reaches the top-5% tier (**Fig. 3c,d**). Consistent with this, the five top-ranked sites of each held-out transcript clustered near the top of its measured activity distribution, most falling within the top 15% (**Fig. 3e**). To visualize ranking quality directly, we compared the measured activity of the sensors built from the top-15 and bottom-15 ranked sites in each held-out transcript. In all six transcripts, top-ranked sites yielded significantly higher-activity sensors than both the bottom-ranked sites and the transcript-wide mean (**Fig. 3f**). Together, these results show that a pairwise model generalizes across transcripts to prioritize high-performing target sites.

To ask whether this ranking ability could have been obtained from existing tools, we compared DeSTAR to Toehold-VISTA^12^, a machine-learning model for target-site selection for conventional toehold switches. Toehold-VISTA was built on a sensor system that differs from Detectron in both switch architecture (25- vs 30-nt toehold, 8- vs 6-bp stem, 33- vs 36-nt target site) (**Extended Data Fig. 4a**) and output modality: conventional toehold switches report through translation so their activity is shaped substantially by translational determinants (RBS accessibility, start-codon context, reporter expression) that have no counterpart in the Detectron architecture, where the readout is a translation-independent, reverse-transcribed DNA barcode. Because the two systems differ in both respects, a model trained on conventional toehold switches cannot be assumed to transfer directly to Detectron.

Consistent with this, Toehold-VISTA applied as published did not recover the Detectron activity ranking. Across all six held-out transcripts, Toehold-VISTA’s top-ranked sites were no more active than either its bottom-ranked sites or the transcript-wide mean: its top-15 and bottom-15 predicted sites were statistically indistinguishable in five of six transcripts, and in DENV its top-predicted sites were significantly less active than its bottom-predicted (**Extended Data Fig. 4b**). Whether this reflects the difference in switch architecture, in output modality, or both remains to be established; regardless, it indicates that target-site selection for Detectron is a distinct problem that benefits from a purpose-built model.

### A bottom-stem pairing grammar is the dominant determinant of sensor performance

To identify which features the model relies on when ranking sites, we retrained the ranking model on all six libraries (10 seeds) and applied TreeSHAP^16^. The 20 features and their five-class grouping are listed in **Extended Data Fig. 5a** and **Supplementary Table 2**. Because the model scores candidates from pairwise feature differences, each SHAP value quantifies how much the difference in a given feature between two candidate sites contributes to the model’s decision of which site ranks higher. We summarized these as the mean absolute SHAP per feature (**Extended Data Fig. 5b)** and per feature class (**Fig. 4a**). Attribution identified the bottom-stem motif as the most-used feature class (∼42% of total |SHAP|), followed by sensor folding ΔG (∼22%) (**Fig. 4a**). Attribution reflects how heavily the model uses a feature, not whether that feature is necessary or sufficient. Removing each feature class in turn showed that deleting the bottom-stem motif class collapsed held-out precision (ΔP@10 = −0.58, Wilcoxon *P* = 0.031), establishing it as the dominant determinant of ranking, whereas deleting the sensor folding ΔG class had no significant effect (ΔP@10 ≈ 0, Wilcoxon *P* = 1.0) despite its being the second most attributed class (**Fig. 4b**). Training on each class in isolation (**Fig. 4c**) told the same story from the opposite direction: no single class approached the full 20-feature model (P@10 = 0.77); the bottom-stem motif was the strongest and most consistent single class (P@10 = 0.30, above chance in 5/6 transcripts), whereas the next-highest, relative transcript position, was powerful in some transcripts (3/6) but uninformative in others. This ranking of class necessity was consistent across precision cut-offs (**Extended Data Fig. 5c**).

**Figure 4.**
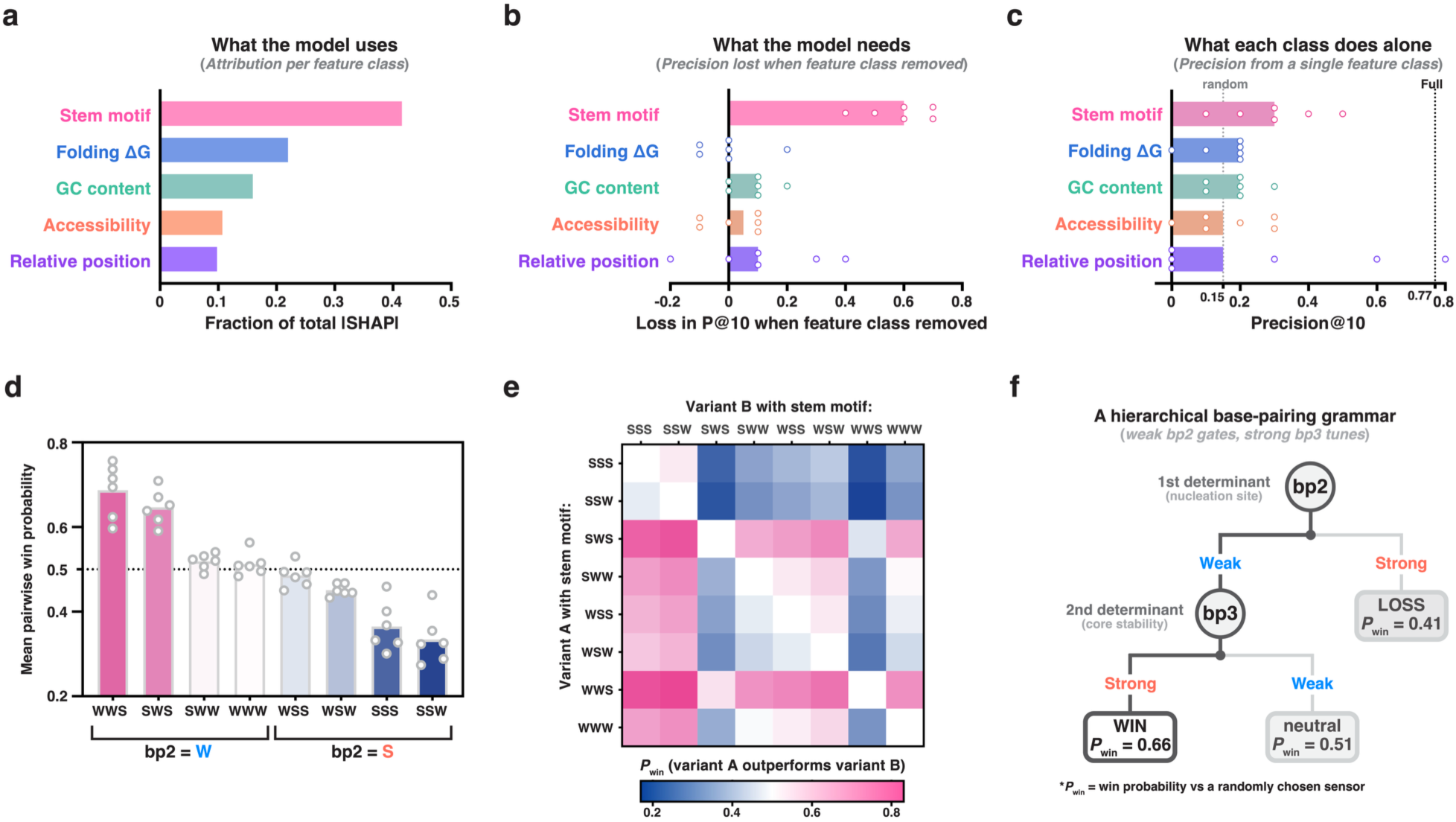
A bottom-stem pairing grammar is the dominant determinant of sensor performance. **a**, What the model uses: mean absolute SHAP (TreeSHAP) per feature class, expressed as a fraction of total |SHAP|; ranking model retrained on all six libraries, averaged over 10 seeds. **b**, What the model needs: loss in precision@10 (responsive, top-15% tier) when each feature class is removed, evaluated by leave-one-transcript-out cross-validation; for each held-out transcript, candidate scores were averaged across the 10 seeds into a single consensus ranking before computing precision. Ablated versus full models were compared across the six held-out transcripts by two-sided Wilcoxon signed-rank test: Bottom-stem motif removed versus full, *P* = 0.031; all other feature classes not significant. *n* = 6 transcripts; bars, mean. **c**, What each class does alone: precision@10 of the ranking model trained on each feature class alone, evaluated by LOTO; dotted lines, random (0.15) and full 20-feature model (0.77). *n* = 6 transcripts; bars, mean. **d**, Mean pairwise win probability for each motif — the probability that a variant carrying that motif has higher measured activity than a randomly chosen variant from the same transcript — grouped by second base-pair identity (bp2 = W versus bp2 = S). Variants with a weak bp2 showed higher activity than those with a strong bp2; two-sided Wilcoxon signed-rank test on the per-transcript difference versus zero: Δmean log₂(ON/OFF) = +0.30, *P* = 0.031. *n* = 6 transcripts. Bars, mean; dotted line, 0.5. **e**, Pairwise win-probability matrix: color indicates probability a variant with the row motif outperforms a variant with the column motif, computed within each transcript and averaged over six libraries. **f**, The motif grammar as a decision rule: the second base pair (bp2) is the primary determinant and, within weak-bp2 sensors, the third base pair (bp3) a secondary determinant. *P*_win_, win probability versus a randomly chosen sensor (within transcript, averaged over six libraries).

The sensor folding ΔG paradox (high attribution but negligible necessity or standalone power) reflects redundancy: the class’s attribution is almost entirely carried by hairpin ΔG feature (∼92% of the class’s |SHAP|), which is 94% reconstructible from the model’s other, non-thermodynamic features (**Extended Data Fig. 5d**), mainly invasion-region GC content (**Extended Data Fig. 5e**). Sensor folding energetics are thus already implicit in the sequence-composition features the model uses, leaving the bottom-stem motif as the single dominant, non-redundant determinant of predicted sensor performance.

We next resolved the bottom-stem motif into its internal grammar. Computing pairwise win probabilities among the eight strong/weak motifs of the three toehold-proximal base pairs (bp1-bp3; **Fig. 2e**) within each transcript revealed a consistent hierarchy (**Fig. 4d,e**). The central base pair (bp2) was the primary determinant: all four weak-bp2 motifs outperformed a randomly chosen sensor (mean pairwise win probability > 0.5) while all four strong-bp2 motifs fell below 0.5 (**Fig. 4d**). Within the winning weak-bp2 motifs, a strong third base pair (bp3) acted as a secondary determinant, so that the weak-base/strong-core motif WWS ranked highest. This hierarchy reduces to a simple rule (**Fig. 4f**): a weak bp2 gates a sensor into the high-performing class, after which a strong bp3 tunes it toward the top. This grammar recapitulates the biophysical architecture read directly from the raw sequences (**Fig. 2d**): the weak second base pair (bp2) that gates switching and the strong core that stabilizes the OFF state (bp3 onward) are the same weak-base/strong-core polarity, now resolved to the single most decisive base pairs.

These two roles have distinct biophysical origins. Strengthening the core (bp3-8) monotonically deepened the computed OFF-state hairpin, expected to make the OFF state more resistant to spontaneous opening, and measured ON/OFF rose in parallel (**Extended Data Fig. 6a**). More informatively, the model left the toehold-terminal pair bp1 essentially unconstrained among the high-performing motifs while strongly favoring a weak bp2, counterintuitive since bp1 lies closer to the invading target. This is consistent with what terminal base-pair fraying predicts^14^. As the terminal pair of the stem, bp1 is dynamically part of the single-stranded toehold junction, opening ∼25% of the time for the free switch and ∼17% even with the target bound to the toehold — an effective free-energy cost of only 0.96 kcal/mol to open, versus 2.57 kcal/mol for bp2 (**Extended Data Fig. 6b,c**). Although bp1 favors pyrimidines in the raw sequence logos (**Fig. 2c**), this preference is near-uniform among high-performing sites and did not improve held-out ranking when encoded explicitly, either as a pyrimidine/purine feature or a full four-base encoding (**Extended Data Fig. 6d**), indicating that bp1 identity does not discriminate among the top-ranked sites. Therefore, bp1’s identity is unlikely to gate switching, and the first reliably paired position, bp2, becomes the first determinant of invasion nucleation. Beyond bp2 the target branch-migrates through the core by replacing each pair with identical strength, an essentially isoenergetic process whose net free energy is insensitive to core composition^17,18^, which leaves bp3-8 free to serve OFF-state stability. Consistent with this view, only sensors combining a weak bp2 with a strong core show the highest mean performance (**Extended Data Fig. 6e**).

The magnitude of this weak-bp2 advantage was itself context-dependent, in a coherent biophysical pattern: across all six transcripts it was largest at the least-accessible (most buried) invasion sites and diminished as accessibility rose, while growing with core strength and local GC content. Each dependence was consistent across all six transcripts (**Extended Data Fig. 7a-c**) and remained significant in a joint model that adjusts for the other two covariates (**Supplementary Table 4**). A weak bp2 thus contributes most where nucleation is otherwise rate-limiting and the OFF state is otherwise well-stabilized, a context-dependence the learned ranker exploits but a fixed motif rule cannot capture.

### DeSTAR prospectively enriches its top-ranked candidates for functional sensors in an unseen transcript

Finally, we asked whether DeSTAR could prospectively select functional sensors for a transcript it had never seen. After training on all six libraries, we froze the model and applied it, without any further tuning or feature selection, to a new target, the SARS-CoV-2 nucleocapsid (N) gene transcript, ranking every candidate site across the transcript (**Fig. 5a**). We then built the ten top-ranked sensors and, as a baseline, twenty sensors targeting randomly selected sites, and measured their activity by two independent formats. In the clonal format, each sensor was assayed individually by qPCR; in the pooled format, all sensors were combined in a single mixed culture and quantified by barcode sequencing (**Fig. 5a**).

**Figure 5.**
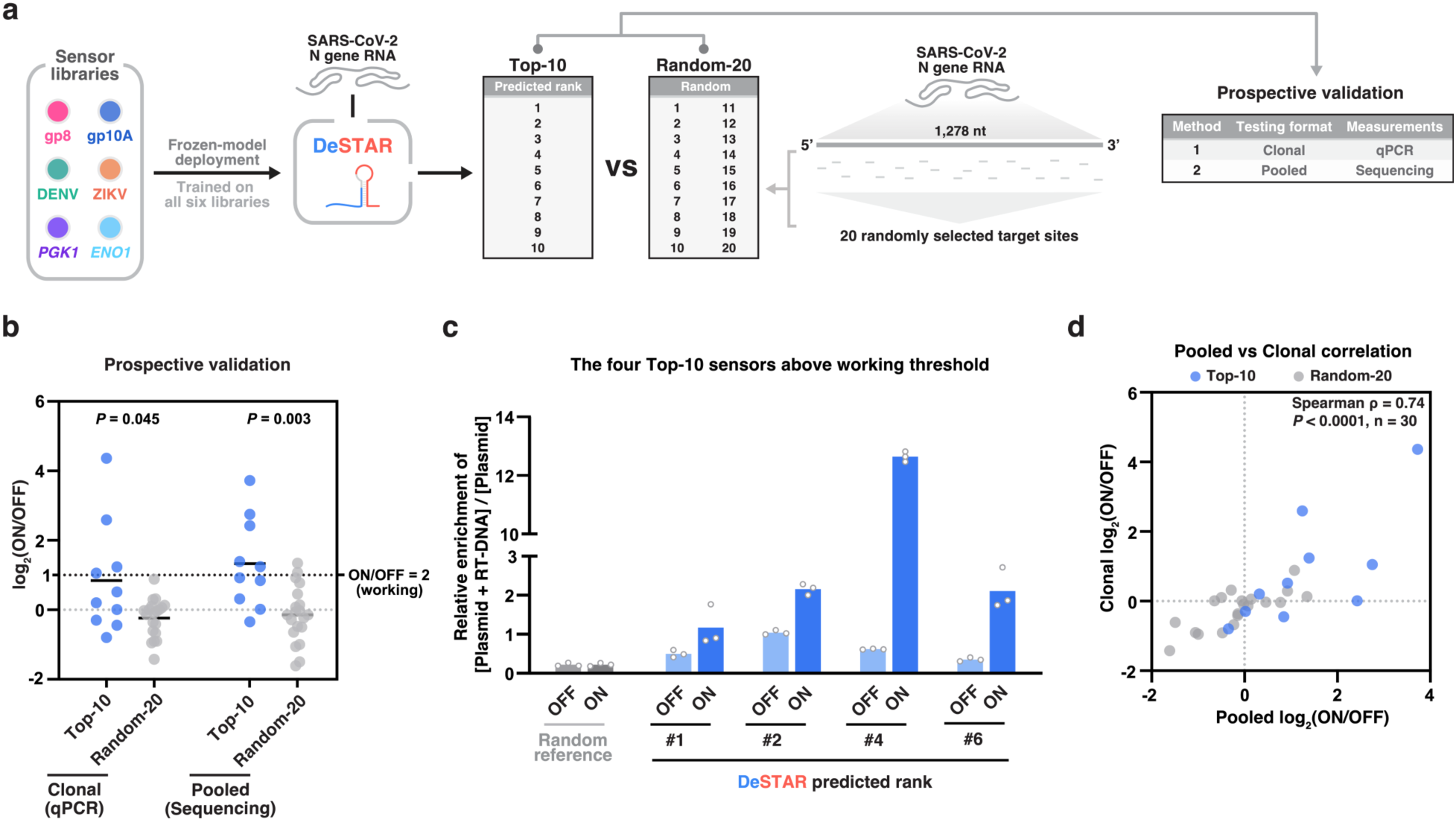
DeSTAR prospectively enriches its top-ranked candidates for functional sensors in an unseen transcript. **a,** Prospective-test design. The final ranking model (DeSTAR), trained and frozen on all six tiling libraries, was deployed and tested on SARS-CoV-2 nucleocapsid (N) gene RNA. All 1,246 overlapping 33-nt candidate target sites were scored and ranked; the 10 top-ranked sites (Top-10) were compared against 20 sites selected at random from the remaining candidates (Random-20). Selected sensors were validated in two independent formats: (1) clonal, with qPCR readout, and (2) pooled, with sequencing readout. **b,** Prospective validation. log_2_(ON/OFF) of Top-10 (blue) versus Random-20 (grey) sensors, measured clonally by qPCR (left) and in a pooled culture by sequencing (right). Each point is one sensor; bars indicate the mean. The dashed line marks the working threshold (ON/OFF = 2). *P* values, two-sided Mann-Whitney *U* test (clonal, *P* = 0.045; pooled, *P* = 0.0030). **c,** The four Top-10 sensors exceeding the working threshold (DeSTAR predicted ranks #1, #2, #4, and #6), assayed clonally by qPCR. Enrichment of RT-DNA + plasmid over plasmid alone is shown in the absence (OFF) or presence (ON) of the target transcript; a representative random-tiling sensor is shown as a reference. Closed circles indicate three biological replicates. Bars represent the mean of three biological replicates. **d,** Concordance between the two readouts. Clonal log_2_(ON/OFF) versus pooled log_2_(ON/OFF) for all 30 sensors (Top-10 blue; Random-20, grey). Spearman rank correlation ρ = 0.74, *P* < 0.0001, *n* = 30; dashed lines mark ON/OFF = 1 on each axis.

By both readouts, the top-ranked sensors were significantly more active than randomly selected sites (Mann-Whitney *P* = 0.045 clonal, *P* = 0.0030 pooled; **Fig. 5b**), and enriched for functional sensors (ON/OFF ≥ 2) over random site selection (4/10 versus 0/20 in the clonal assay and 5/10 versus 2/20 in the pooled assay). For the four top-10 sensors that cleared the working threshold, the ON signal was induced well above the OFF baseline, reaching ∼21-fold (ON/OFF = 20.6) for sensor #4, whereas the non-responsive random reference showed no ON/OFF separation (**Fig. 5c**). Critically, the two readouts, which differ in both testing format and measurement method, agreed: per-sensor activity was strongly correlated between clonal qPCR and pooled sequencing (Spearman ρ = 0.74, *P* < 0.0001, *n* = 30), showing that the ranking holds across independent experimental platforms (**Fig. 5d**). Together, these results show that DeSTAR, frozen and applied prospectively, enriches its top-ranked candidates for functional sensors in a transcript absent from its training, reducing the empirical screening needed to build a working Detectron sensor for a new target.

## DISCUSSION

Efforts to improve toehold switch sensors have concentrated on the switch itself — forward-engineered^1^ and, more recently, deep-learning-based designs^10,11^ — yet the site within a transcript a sensor targets has received far less attention, despite being a major determinant of performance in its own right. This gap is concrete for Detectron^13^, a recently introduced RNA sensor that reports target recognition as a reverse-transcribed DNA barcode rather than a translated protein: its switch architecture was optimized by high-throughput screening, but how to select the target site within a transcript was not addressed, and no target-site model has existed for it. The one machine-learning tool for target-site selection, Toehold-VISTA^12^, was developed for translation-based switches and performs well in that setting, but its rankings did not predict Detectron activity across our six transcripts (**Extended Data Fig. 4**); whether this reflects the different output modality, differences in switch architecture, or both remains to be determined. We therefore built DeSTAR, a pairwise ranking model trained directly on Detectron activity across six tiled transcripts, and show that, frozen and applied prospectively, it enriches its top-ranked candidate sites for functional sensors in a transcript it had never seen.

One defining feature of our approach is its resolution. Sensor activity varies at single-nucleotide resolution: target sites one nucleotide apart differ in activity nearly as much as randomly chosen ones (**Extended Data Fig. 1**), so there is no broadly favorable region to target and each candidate site must be evaluated individually by the model. Tiling and scoring at single-nucleotide resolution were therefore essential — coarser sampling would skip over functional sites, and the best site for a given transcript can fall between more widely spaced positions.

Beyond prediction, the translation-independent readout clarified the sequence rules of activation. Because Detectron reflects strand invasion directly, without the translational layer that conventional reporters impose, we could resolve the toehold-proximal stem base pair by base pair and assign distinct roles: a weak second pair (bp2) gates activation, a strong third pair (bp3) tunes it, and the terminal pair (bp1) frays without gating (**Fig. 4**, **Extended Data Fig. 6**). This positional grammar, rather than the stem’s overall G·C content, is the dominant determinant of performance (**Fig. 4**), and it explains the single-nucleotide sensitivity: a one-nucleotide shift rewrites which base pair occupies each stem position, whereas the site’s accessibility and overall base composition change little. Consistent with this, the sites that yield high-performing sensors are interspersed rather than clustered, as they would be if regional accessibility were the primary driver.

Accessibility and the broader structure of the target within the full-length transcript instead play a secondary, tuning role and set how strongly the grammar applies — the weak-bp2 advantage is largest where the invasion site is otherwise buried and the core otherwise stable (**Extended Data Fig. 7**) — so sites are best assessed within the full transcript rather than as isolated short triggers, consistent with the importance of transcript context also noted for translation-based switches^12^. The learned ranker captures this positional, context-dependent grammar directly from sequence, recapitulating the biophysics of a weak pair that nucleates invasion over a strong core that stabilizes the OFF state (**Fig. 2**). These features also make Detectron a scalable platform for dissecting the sequence determinants of strand invasion: because its DNA-barcode output couples the invasion event to a readout without a translational intermediary and can be read for thousands of variants in parallel by sequencing, the mechanism can be probed systematically at single-nucleotide resolution.

Applied to the SARS-CoV-2 nucleocapsid (N) transcript without further tuning, DeSTAR concentrated functional sensors into its top-ranked candidates: its top-10 were significantly more active than randomly selected sites and enriched for working sensors (ON/OFF ≥ 2) in both a clonal qPCR assay and an independent pooled-sequencing assay, which agreed closely (**Fig. 5**). For a user building sensors against a new transcript, this converts a blind empirical screen into a short, prioritized list, reducing the number of constructs that must be tested to obtain a working sensor.

The principal limitation is the breadth of our training data. The model is trained and evaluated on six transcripts, and functional sensors are rare even in the best of them, so its performance is bounded more by this breadth and by the rarity of functional sites than by per-transcript depth; additional tiled transcript libraries are the clearest path to a stronger ranker. For the same reason, DeSTAR ranks sites within a transcript but cannot predict a transcript’s overall yield of functional sensors, which varies widely across targets — such a transcript-level prediction would require training across many transcripts, so we cannot yet flag, before testing, that a target is intrinsically low-yield. A calibrated low-yield warning is a natural direction as more transcripts are profiled. Our determinants are also inferred from naturally occurring target sequences sampled by tiling, rather than from a library that systematically varies sensor composition: we have not surveyed the full combinatorial space of the eight stem base pairs, and some determinants may be entangled with correlated features of the tiled sequences. A purpose-built library that independently varies stem composition against cognate sites embedded in full-length transcripts would be needed to establish these rules causally and to resolve the detailed physical mechanism of strand invasion that underlies them.

More broadly, DeSTAR shows that target-site selection for a DNA-output RNA sensor is a learnable problem, distinct from the translation-based rules that preceded it, and that a model trained on one sensor chemistry can be frozen and deployed prospectively on unseen transcripts. As more Detectron libraries accumulate, the same framework should sharpen and could be extended to reliable, per-target estimates of expected yield and to related RNA-sensing platforms^19,20^ — making rational, first-pass sensor design practical for arbitrary RNA targets.

## METHODS

All biological replicates were taken from distinct samples, not the same sample measured repeatedly. For Detectron tiling libraries, each biological replicate is an independent electroporation and expression of the libraries into the strain *E. cloni* 10G. All statistical tests and *P*-values are included in Supplementary Table 4.

### Plasmid construction and architecture

Detectron constructs were cloned into an ampicillin-resistant pBR322-derived plasmid containing a ColE1/pMB1-family origin together with bom and rop. The retron ncRNA (msr-msd) and Eco1 reverse transcriptase were expressed as a polycistronic cassette under constitutive *glpT* promoter and terminated using B0014 terminator. Trigger RNAs (or target RNAs) were encoded on the same plasmid under the indicated promoters (J23100 or pBAD) and terminated using the T7 transcription terminator (T7Te). Detailed construct annotations for all key constructs are provided in Supplementary Table 5.

### qPCR

RT-DNA was quantified by qPCR as described previously^13,21^, using two primer sets: Eco1_wt_2F/Eco1_wt_R and Eco1_wt_3F/Eco1_wt_R. The first pair anneals within the msd region and therefore amplifies both plasmid DNA and RT-DNA, whereas the second pair targets a region outside the msd and amplifies plasmid DNA alone. For each biological replicate, the difference in cycle threshold (ΔCt) between the two primer sets was calculated, and the enrichment of the combined [Plasmid + RT-DNA] signal relative to the [plasmid] signal within each sample was expressed as 2^-ΔCt^. Primer sequences are provided in Supplementary Table 6.

### Barcode design

Each target site was assigned three 13-nt DNA barcodes designed with a minimum pairwise Hamming distance of 4 to tolerate sequencing errors and support reliable demultiplexing, and with GC content restricted to 30–70% to limit secondary structure and ensure uniform amplification.

### Detectron tiling library cloning

Detectron tiling libraries were constructed essentially as described previously^13^, with the following modification: the tiling libraries were assembled in a single SapI Golden Gate step rather than the two-step SapI/BsaI strategy used previously. Briefly, synthesized oligo pools (Twist Bioscience) were amplified with skpp15 primer and assembled into the Detectron backbone using the SapI Type IIS restriction enzyme. For each transcript, two libraries were constructed: an ON library, constitutively expressing trigger RNA, and an OFF library, lacking trigger RNA expression. Cleaned-up Golden Gate reactions (6 ul) were mixed with 25 ul of *E*. *cloni* 10G cells (LGC Biosearch Technologies, COMCEL-018) were added to 0.1-cm gap electroporation cuvettes (BioRad, 1652083) and electroporated at 1,800 V, 25 μF, and 200 Ω. Cells were immediately recovered in 1 mL of pre-warmed recovery medium and shaken for 1 h at 37 °C. Serial dilutions were plated on LB supplemented with carbenicillin. CFUs were counted the following day, and libraries were carried forward only when the estimated coverage exceeded 200×.

### Library assay and sequencing

Assay and quantification followed the Detectron pipeline^13^, with minor modifications. Briefly, each library was prepared as paired ON and OFF sublibraries (cognate target transcript present or absent, respectively) and introduced into *E. cloni* 10G cells by electroporation, with three independent electroporations per sublibrary serving as biological replicates. After overnight growth, barcode-encoded RT-DNA and plasmid DNA were co-extracted using the QIAGEN Plasmid Plus Midi Kit; an aliquot of the eluate was reserved for plasmid quantification, and the remainder was enriched for single-stranded RT-DNA by plasmid depletion (Zymo ssDNA Clean & Concentrator Kit). The RT-DNA and plasmid fractions were separately indexed by two rounds of PCR to add sequencing adapters and sample indexes as described previously^21–23^, and sequenced on an Oxford Nanopore PromethION. Sequencing data were collected using MinKNOW version 25.09.16 (Bream version 8.8.3, Configuration version 6.8.9 and MinKNOW Core version 6.8.11) and basecalled with Dorado version 7.11.2 FASTQ file were demultiplexed and processed using custom code (https://github.com/Shipman-Lab/DeSTAR). For each variant, RT-DNA abundance was normalized to the abundance of its cognate plasmid, sequenced in parallel, to yield per-variant activity. All primer sequences are provided in Supplementary Table 6.

### Single-nucleotide sensitivity analysis

For each transcript, we computed the absolute difference in log_2_(Average ON/OFF) between variants targeting adjacent (one-nucleotide-apart) sites and between an equal number of randomly drawn variant pairs. To test whether adjacent differences exceeded measurement noise, we defined a per-pair noise floor — the median absolute difference expected from replicate noise alone — as 0.6745 × √(SE*ᵢ*² + SE*ⱼ*²), where SE is the standard deviation of variant’s Dixon-retained replicate log_2_(ON/OFF) divided by √*n*. Significance was assessed by a one-sided sign test (exact binomial) of whether adjacent |Δlog₂(ON/OFF)| exceeds its pair-specific noise median more often than chance (0.5), computed per transcript and pooled (**Supplementary Table 4**).

### Sequence logo analysis

To visualize sequence features associated with sensor performance, enrichment sequence logos were generated with Logomaker^24^ over the 33-nt switch region (25-nt single-stranded toehold and 8-nt variable stem1). Within each tiling library, candidate sensors were ranked by log_2_(ON/OFF) and partitioned into a high-performing group (top 15%) and a low-performing group (bottom 15%). For the pooled logo, the top-15% and bottom-15% sequences were combined across all six libraries; per-transcript logos were computed within each library and plotted on a common symmetric y-axis for cross-transcript comparison. At each position, A/C/G/U counts were tallied per group with a pseudocount of 0.5 and normalized to per-position frequencies. Letter heights represent the per-base enrichment, log_2_(high-group frequency / low-group frequency): bases enriched in high-performing sensors are drawn above the axis and depleted bases below it. Logos were rendered with logomaker v0.8.7 (Python 3.11).

### Machine-learning target-site ranking model

#### Data and labels

Six single-nucleotide-resolution tiling libraries (gp8, *n* = 1,345; gp10A, *n* = 1,006; DENV, *n* = 1,453; ZIKV, *n* = 1,174; *PGK1*, *n* = 1,135; *ENO1*, *n* = 1,270; 7,383 target-site variants total) were used to train and evaluate the model. Each variant’s activity was taken as its replicate-averaged ON/OFF ratio and transformed to log_2_(ON/OFF). The appended control region of each non-Zika library was excluded from all training and evaluation.

#### Features

Each candidate target site was represented by 20 features spanning five classes: (i) stem motif — eight one-hot indicators of the bottom-three-base-pair stem strong/weak motif (SSS, SSW, SWS, SWW, WSS, WSW, WWS, WWW); (ii) folding ΔG — two raw sensor-folding free energies (hairpin ΔG and toehold-hairpin ΔG); (iii) GC content — four transcript-relative GC content features (target-window, seed-region, invasion-region and local context GC); (iv) accessibility — five transcript-relative RNA accessibility features (mean and minimum single-base accessibility over the 33-nt target window, and full-window, seed-region and invasion-region accessibility); and (v) relative transcript position. Because the model is tree-based and operates on feature differences, features were not standardized; missing values were median-imputed within a scikit-learn Pipeline^25^. These arose only for the 5’ terminal sites of gp8 and gp10A (expressed without flanking), where accessibility is undefined; because the imputation contributes a neutral, zero-difference term, excluding these sites left held-out precision @5/10/15 unchanged and placed no terminal site in the model’s top-15.

#### Pairwise learning-to-rank formulation

The task was cast as pairwise ranking. Within each transcript, variant pairs (A, B) were formed and labelled *L* = 1 if log_2_(ON/OFF)A > log_2_(ON/OFF)B and 0 otherwise, with feature vector ΔX = X_A_ – X_B_. Each sampled pair was added symmetrically — both (ΔX, *L*) and (–ΔX, 1 – L) — so the classifier learns an antisymmetric win-probability function *P*(A > B). Pairs were built independently within each transcript and never across transcripts, preserving cross-transcript independence. A gradient-boosted decision-tree classifier (XGBoost; n_estimators = 500, max_depth = 5, learning_rate = 0.05, subsample = 0.9, colsample_bytree = 0.9, reg_lambda = 1.0) was trained on the pooled pairwise examples from the training transcripts.

#### Top-focused pair sampling and zone weighting

To concentrate the model’s discriminative capacity on high-performing sites, each variant was assigned to a disjoint within-transcript performance band by activity percentile — top 5%, 5-10%, 10-15%, or the remaining bottom 85% — and up to 15,000 training pairs per transcript were drawn from three combined pools: (i) a nested, band-stratified pool that preferentially samples cross-band comparisons among the top bands (90% of the budget); (ii) a top-focused pool pairing top-ranked variants against the rest (top fraction 0.125, “one-in-top” model); and (iii) random pairs filling the remainder. All pairs required a minimum activity separation of |Δlog_2_(ON/OFF)| ≥ 0.10. Pairs spanning the top bands were then up-weighted in the training loss via per-sample weights (top-5% vs 5-10%: ×2.5; top-5% vs 10-15%: ×2.0; 5-10% vs 10-15%: ×1.2; all other pairs ×1.0).

#### Scoring and cross-validation

A held-out transcript’s candidate sites were ranked by pairwise-win score: each candidate received the mean predicted win probability *P*(i > j) against all other candidates in the same transcript. The model was evaluated by leave-one-transcript-out cross-validation — trained on five libraries and used to rank the held-out sixth, repeated for all six — with no information from the held-out transcript entering feature construction, imputation, pair building or training. The entire procedure was repeated over 10 random seeds and averaged.

#### Performance metrics

Within each held-out transcript, sites were assigned to performance tiers by within-transcript quantile: responsive (top 15%), strong (top 10%), and elite (top 5%; tiers required ≥ 10 positives). For each tier we report precision@*k* (P@5/10/15), the fraction of the model’s top-k ranked sites falling in the tier, and the enrichment factor EF@*k* = P@*k* ÷ tier base rate (0.15, 0.10, 0.05), i.e. fold-enrichment of the top-k picks over random tiling.

### Feature-class importance analysis

To interpret the ranking model, the 20 features were grouped into the five classes defined above — stem motif, folding ΔG, GC content, accessibility and relative transcript position — and three complementary analyses were performed on the pairwise ranking model under leave-one-transcript-out (LOTO) cross-validation using the final model configuration described above, averaged over 10 seeds.

#### Attribution — what the model uses

For each LOTO fold and seed, TreeSHAP^16^ (shap.TreeExplainer) was applied to the trained XGBoost pairwise classifier. Because the model operates on feature differences (X_A_ – X_B_), each SHAP value quantifies how a feature difference contributes to the predicted win probability. The mean absolute SHAP value was computed per feature, summed within each feature class, and expressed as a fraction of the total mean absolute SHAP across all features; class fractions were averaged across folds and seeds.

#### Necessity — what the model needs

For each feature class, the model was retrained with that class’s features removed (using only the remaining four classes) and re-evaluated by full LOTO. The importance of a class was quantified as the loss in precision@10 relative to the full 20-feature model (full P@10 – ablated P@10). Significance was assessed by Wilcoxon signed-rank test comparing ablated versus full-model P@10 across the paired seed × held-out-transcript observations.

#### Sufficiency — what each class does alone

For each feature class, a model was trained using only that class’s features and evaluated by full LOTO, yielding precision@10 for each class in isolation. Single-class precision was compared against the random-tiling base rate (0.15) by Wilcoxon signed-rank test; the full 20-feature model (P@10 = 0.77) is shown as the upper reference.

### Bottom-3-bp stem-motif grammar analysis

To characterize the empirical relationship between the bottom-three-base-pair stem motif and sensor performance, each sensor was assigned its stem motif (one of eight strong/weak combinations, SSS-WWW) and pairwise win rates were computed directly from measured activities (log_2_(ON/OFF)), independently of the ranking model.

#### Pairwise win-rate matrix

For each ordered pair of motifs (A, B), the empirical win probability *P*(A > B) was defined as the fraction of all cross-motif sensor pairs — one sensor of motif A, one of motif B — in which the motif A sensor had the higher log_2_(ON/OFF), with ties counted as 0.5 (equivalent to the Mann-Whitney U / AUC statistic). The 8 × 8 matrix was computed within each library and averaged across the six libraries.

#### Mean pairwise win probability

For each motif, a summary score was computed as the mean of its win probabilities against all eight motifs (the row mean of the matrix); a value of 0.5 indicates no net advantage. Motifs were grouped by their second base pair (bp2 = weak vs strong) to display the bp2 effect.

### Prospective validation on an unseen transcript

As a prospective test, the ranking model was frozen and applied to a transcript held out of all model development — the SARS-CoV-2 nucleocapsid (N) gene RNA.

#### Candidate ranking by the frozen model

The final pairwise ranking model was trained on all six tiling libraries using the frozen configuration, with no further tuning. The SARS-CoV-2 N gene sequence was tiled into all overlapping 33-nt candidate target sites (*n* = 1,246), and the final 20-feature set was computed for each site exactly as for the training libraries, with secondary-structure and accessibility features generated using ViennaRNA Package v2.7.0^26^ (RNAfold and RNAplfold^27^) and GC/accessibility features expressed as transcript-relative percentiles. Each candidate was scored by the frozen model (mean pairwise-win score, averaged across the 10 seeds) and ranked.

#### Sensor selection and construction

The 10 top-ranked candidate sites and 20 sites drawn at random from the remaining candidates were selected (30 sensors total), each assigned a unique 13-nt barcode. Sensors were synthesized as gBlocks (IDT) and cloned into the Detectron backbone. For each sensor, activity (ON/OFF) was defined by the plasmid-normalized RT-DNA abundance measured in the presence (ON) versus absence (OFF) of the cognate SARS-CoV-2 N trigger RNA. The prospective comparison evaluated the model’s top-10 predictions against the 20 randomly selected sites.

## ACKNOWLEDGEMENTS

Work was supported by funding from the Bachrach Family Foundation, the National Science Foundation (MCB 2509382), and the National Institute of Allergy and Infectious Diseases (P01AI195327). S.L.S. is a Chan Zuckerberg Biohub – San Francisco Investigator.

## AUTHOR CONTRIBUTIONS

J.H.: Conceptualization, Investigation, Methodology, Visualization, Writing – original draft. S.L.S.: Supervision, Project administration, Funding acquisition, Writing – review & editing.

## COMPETING INTERESTS

J.H. and S.L.S. are named inventors on U.S. Provisional Patent Application No. 63/900,194 covering the Detectron RNA-sensing platform on which this work builds.

## DATA AVAILABILITY

Sequencing data associated with this study are available in the NCBI Sequence Read Archive (BioProject PRJNA1529981). Complete variant, plasmid, primer and statistical summary information are provided in the Supplementary Tables.

## CODE AVAILABILITY

All custom code used to analyze the sequencing data and to train, evaluate, and apply the DeSTAR target-site ranking model is available through the Shipman Lab GitHub repository (https://github.com/Shipman-Lab/DeSTAR). The repository includes scripts to reproduce the main and Extended Data figures from the deposited data, and a tool to rank candidate target sites on new transcripts.

## REFERENCES

1. Green, A. A., Silver, P. A., Collins, J. J. & Yin, P. Toehold switches: de-novo-designed regulators of gene expression. Cell 159, 925–939 (2014).

2. Green, A. A. et al. Complex cellular logic computation using ribocomputing devices. Nature 548, 117–121 (2017).

3. Pardee, K. et al. Rapid, low-cost detection of Zika virus using programmable biomolecular components. Cell 165, 1255–1266 (2016).

4. Takahashi, M. K. et al. A low-cost paper-based synthetic biology platform for analyzing gut microbiota and host biomarkers. Nat. Commun. 9, 3347 (2018).

5. Yan, Z. et al. Programmable fluorescent aptamer-based RNA switches for rapid identification of point mutations. Nat. Chem. 17, 1826–1838 (2025).

6. Yan, Z. et al. Rapid, multiplexed, and enzyme-free nucleic acid detection using programmable aptamer-based RNA switches. Chem 10, 2220–2244 (2024).

7. Hunt, J. P. et al. Towards detection of SARS-CoV-2 RNA in human saliva: a paper-based cell-free toehold switch biosensor with a visual bioluminescent output. N. Biotechnol. 66, 53–60 (2022).

8. Park, S. & Lee, J. W. Detection of coronaviruses using RNA toehold switch sensors. Int. J. Mol. Sci. 22, 1772 (2021).

9. Wessels, H. H. et al. Prediction of on-target and off-target activity of CRISPR–Cas13d guide RNAs using deep learning. Nat. Biotechnol. 42, 628–637 (2023).

10. Valeri, J. A. et al. Sequence-to-function deep learning frameworks for engineered riboregulators. Nat. Commun. 11, 5058 (2020).

11. Angenent-Mari, N. M., Garruss, A. S., Soenksen, L. R., Church, G. & Collins, J. J. A deep learning approach to programmable RNA switches. Nat. Commun. 11, 5057 (2020).

12. Robson, J. M. & Green, A. A. Toehold-VISTA: a machine learning approach to decipher programmable RNA sensor-target interactions. Nucleic Acids Res. 54, gkag097 (2026).

13. Han, J. & Shipman, S. L. Detectrons convert transient RNA sequences into stable DNA barcodes for high-throughput analysis of RNA-dependent processes. Nat. Biotechnol. 10.1038/s41587-026-03334-7 (2026).

14. Nonin, S., Leroy, J.-L. & Guéron, M. Terminal base pairs of oligodeoxynucleotides: imino proton exchange and fraying. Biochemistry 34, 10652–10659 (1995).

15. Chen, T. & Guestrin, C. XGBoost: a scalable tree boosting system. in Proceedings of the 22nd ACM SIGKDD International Conference on Knowledge Discovery and Data Mining 785–794 (ACM, 2016).

16. Lundberg, S. M. et al. From local explanations to global understanding with explainable AI for trees. Nat. Mach. Intell. 2, 56–67 (2020).

17. Srinivas, N. et al. On the biophysics and kinetics of toehold-mediated DNA strand displacement. Nucleic Acids Res. 41, 10641–10658 (2013).

18. Zhang, D. Y. & Winfree, E. Control of DNA strand displacement kinetics using toehold exchange. J. Am. Chem. Soc. 131, 17303–17314 (2009).

19. Kaseniit, K. E. et al. Modular, programmable RNA sensing using ADAR editing in living cells. Nat. Biotechnol. 41, 482–487 (2023).

20. Gootenberg, J. S. et al. Nucleic acid detection with CRISPR-Cas13a/C2c2. Science 356, 438–442 (2017).

21. Lopez, S. C., Crawford, K. D., Lear, S. K., Bhattarai-Kline, S. & Shipman, S. L. Precise genome editing across kingdoms of life using retron-derived DNA. Nat. Chem. Biol. 18, 199–206 (2022).

22. Crawford, K. D., Khan, A. G., Lopez, S. C., Goodarzi, H. & Shipman, S. L. High-throughput variant libraries and machine learning yield design rules for retron gene editors. Nucleic Acids Res. 53, gkae1199 (2025).

23. Khan, A. G. et al. An experimental census of retrons for DNA production and genome editing. Nat. Biotechnol. 43, 914–922 (2025).

24. Tareen, A. & Kinney, J. B. Logomaker: beautiful sequence logos in Python. Bioinformatics 36, 2272–2274 (2020).

25. Pedregosa, F. et al. Scikit-learn: machine learning in Python. J. Mach. Learn. Res. 12, 2825–2830 (2011).

26. Lorenz, R. et al. ViennaRNA Package 2.0. Algorithms Mol. Biol. 6, 26 (2011).

27. Bernhart, S. H., Hofacker, I. L. & Stadler, P. F. Local RNA base pairing probabilities in large sequences. Bioinformatics 22, 614–615 (2006).

