## Extended Data Figures for "High-throughput variant libraries and machine learning enable robust target-site selection for toehold-switch design in Detectrons"

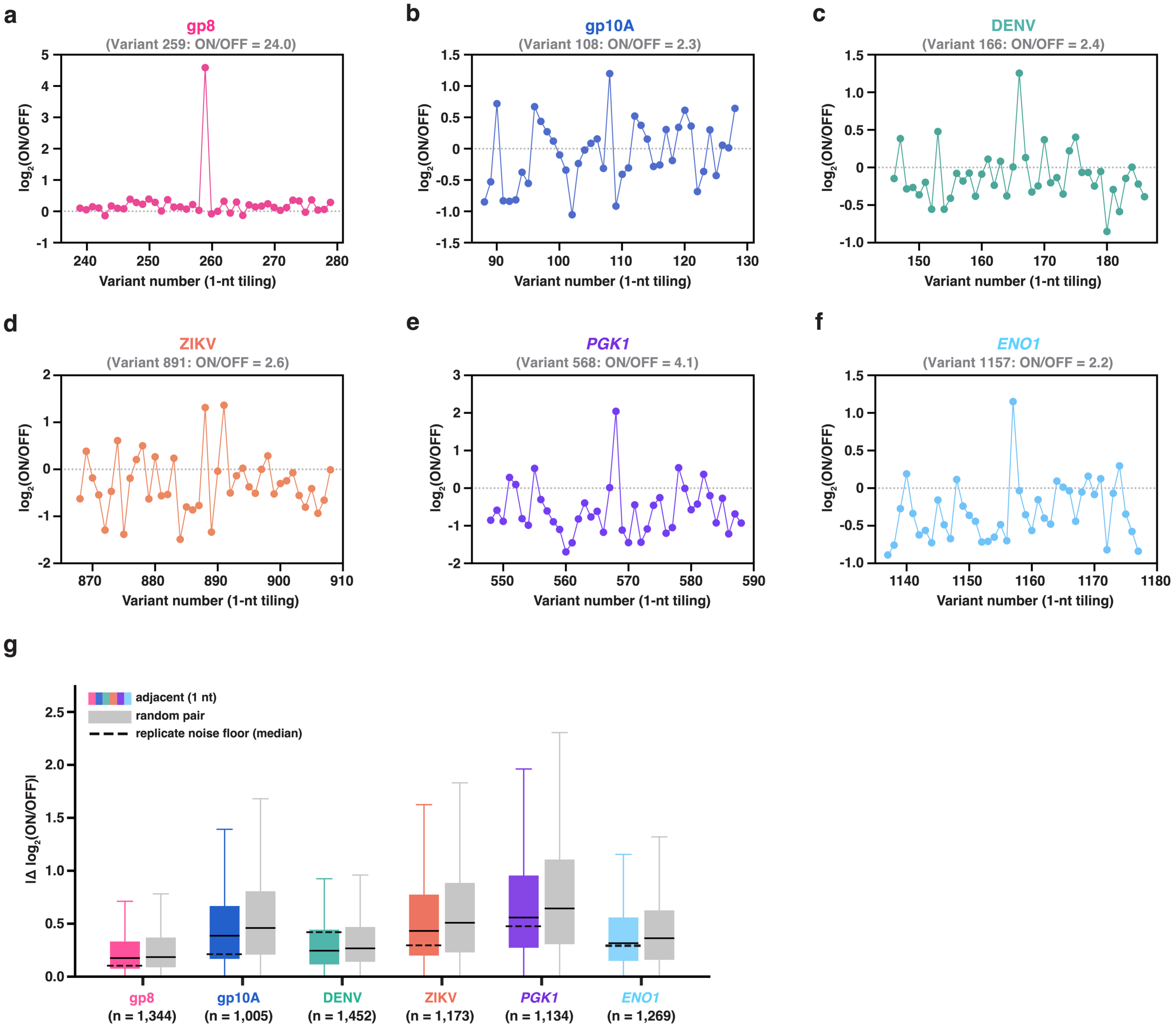


**Extended Data Fig. 1 | Sensor activity varies with target-site position at single-nucleotide resolution.** **a-f**, Representative single-nucleotide tiling windows for each transcript (**a**, gp8; **b**, gp10A; **c**, DENV; **d**, ZIKV; **e**, *PGK1*; **f**, *ENO1*). Each point is one target-site variant — a 33-nt window tiled at single-nucleotide resolution, so consecutive variant numbers differ by a one-nucleotide shift. Each panel shows a 41-variant window centered on a high-activity site; the highlighted variant (variant number and its ON/OFF ratio given in each panel) is the most active site within the window. Activity is $\log_2$(ON/OFF); the dashed grey line marks ON/OFF = 1. In every transcript, a single-nucleotide shift can change a sensor from high-performing to inactive. **g**, Absolute difference in $\log_2$(ON/OFF) between variants whose target sites are one nucleotide apart (adjacent, 1 nt; colored) and between an equal number of randomly chosen variant pairs (random pair; grey), for each transcript. Box, median and interquartile range; whiskers, 1.5× interquartile range; outliers not shown. *n*, number of adjacent 1-nt pairs (below each transcript). Dashed line, median replicate measurement-noise floor — the difference expected between two sites from assay noise alone, had their true activities been equal (Methods). Adjacent-site differences significantly exceeded the noise floor in five of six transcripts (all except DENV; one-sided sign test; per-transcript and pooled statistics in Supplementary Table 4).

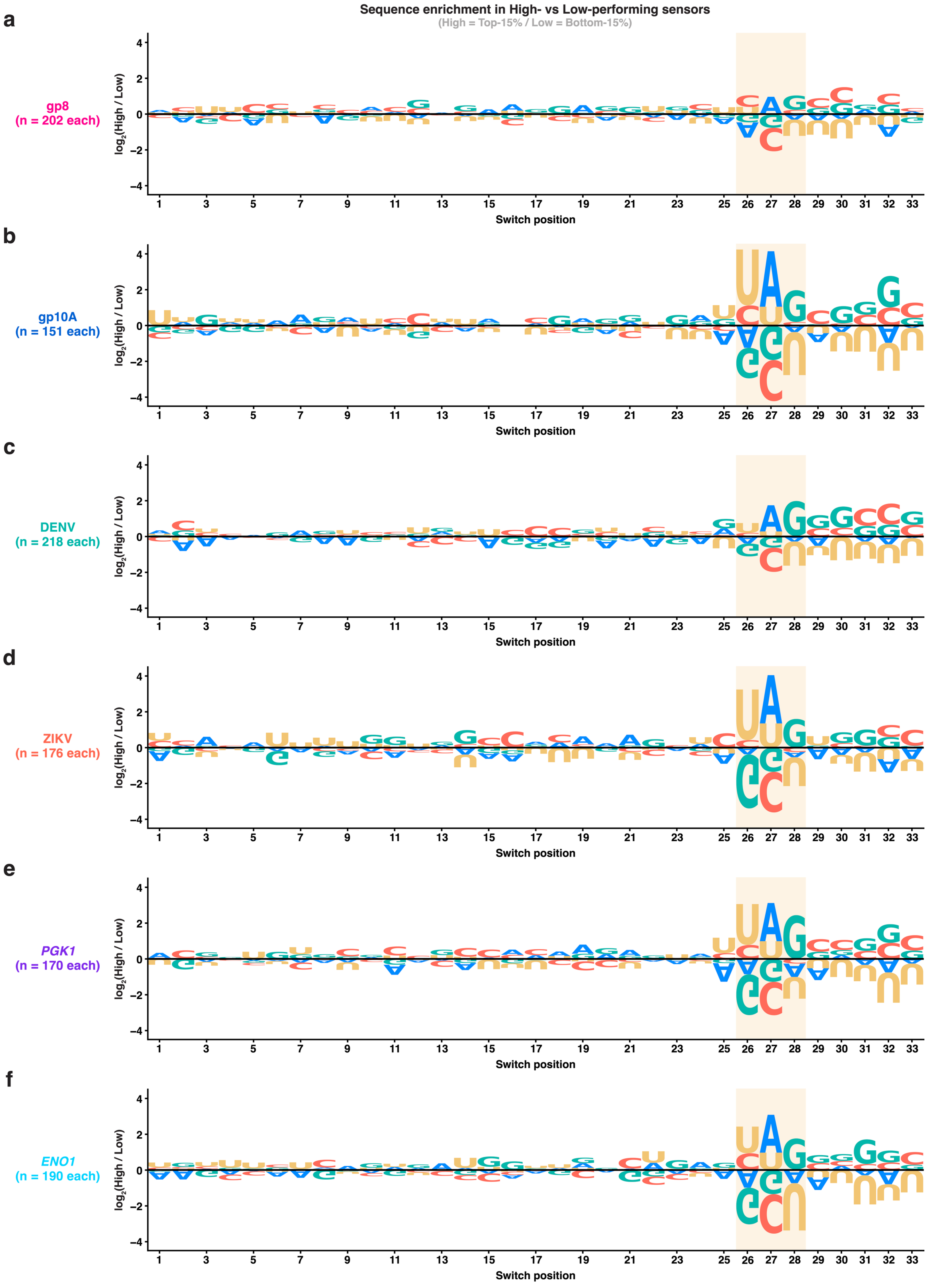


**Extended Data Fig. 2 | The bottom-stem grammar is consistent across all six libraries. a-f**, Per-position sequence enrichment between high- and low-performing sensors, computed separately for each library: **a**, gp8; **b**, gp10A; **c**, DENV; **d**, ZIKV; **e**, *PGK1*; **f**, *ENO1*. At each switch position (toehold, 1-25; stem pairs (bp1-bp8), 26-33), letter height is the $\log_2$ ratio of that nucleotide's frequency in high- versus low-performing sensors (High, top 15%; Low, bottom 15%, ranked within each library by measured $\log_2$(ON/OFF); equal numbers per group, n as indicated in each panel). Letters above the axis are enriched in high performers, below the axis in low performers; nucleotides colored A (blue), C (orange), G (green), U (yellow). Shaded yellow band, bottom-stem positions 26-28 (bp1-bp3), where enrichment concentrates: a weak pair (A/U) is favored at bp2 (position 27) and strong pairs (G/C) through the core (bp3-bp8, positions 28-33). The same pattern recurs in every library despite their differing dynamic range.

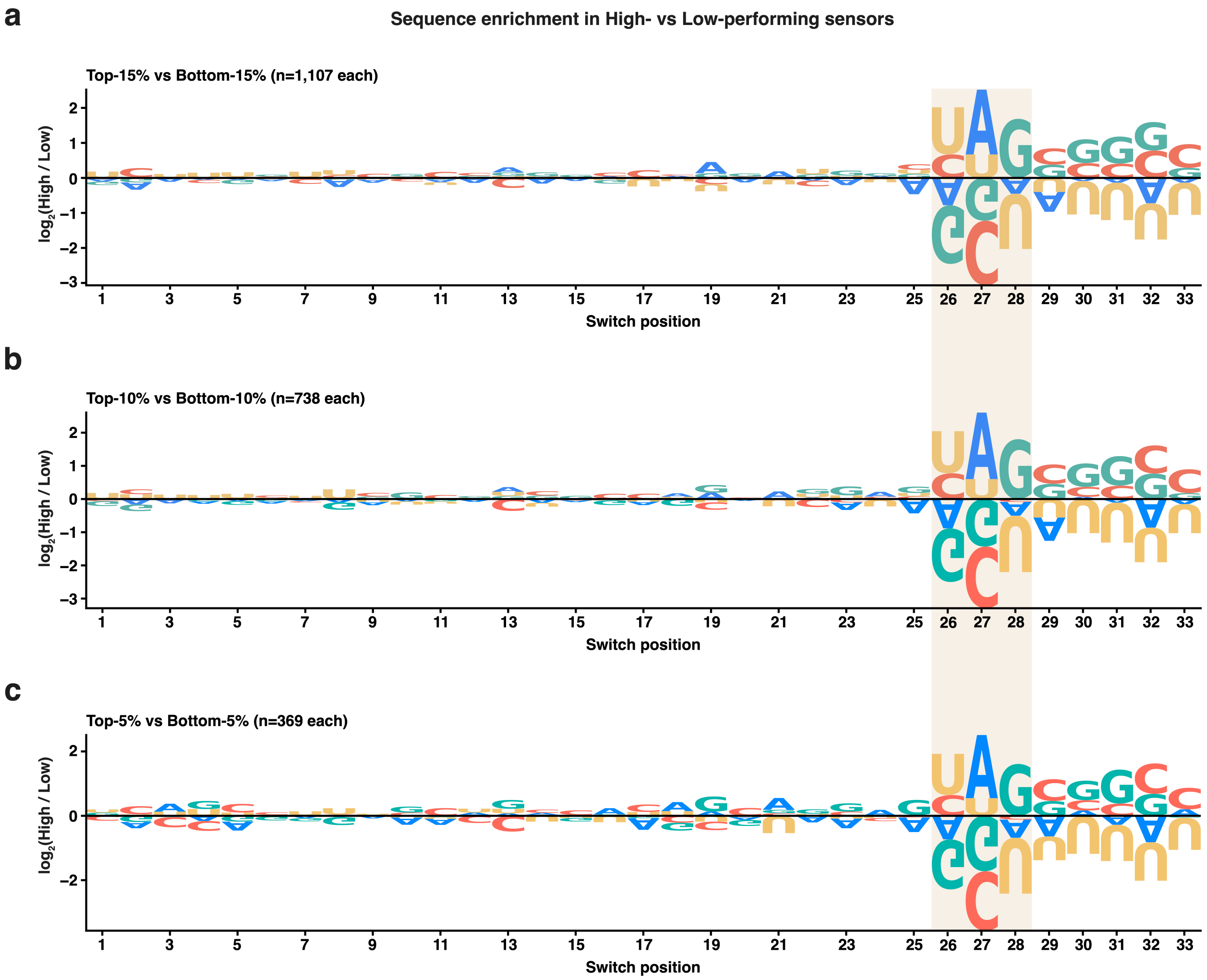


**Extended Data Fig. 3 | The bottom-stem grammar is consistent across performance-tier definitions (all libraries pooled). a-c**, Sequence enrichment between high- and low-performing sensors, pooled across all six libraries, at three increasingly stringent tier definitions: **a**, top versus bottom 15% (*n* = 1,107 each); **b**, 10% (*n* = 738 each); **c**, 5% (*n* = 369 each). Sensors were ranked within each library by measured $\log_2$(ON/OFF), then pooled. Letters above the axis are enriched in high performers, below the axis in low performers; nucleotides colored A (blue), C (orange), G (green), U (yellow). Shaded yellow band, bottom-stem positions 26-28 (bp1-bp3).

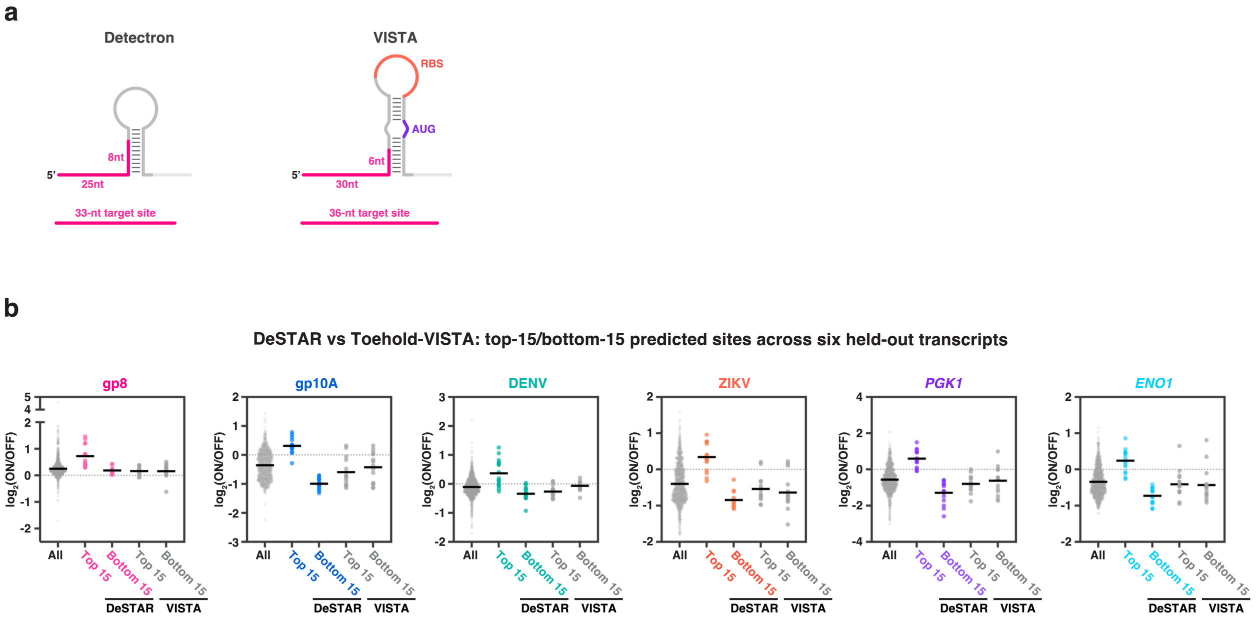


**Extended Data Fig. 4 | DeSTAR, but not an existing protein-output toehold-switch model, predicts DNA barcode-sensor activity. a**, Architectures of the two sensor formats whose target-site rankings are compared. The Detectron sensor (this study; DNA-barcode output) uses a 33-nt target site, comprising a 25-nt toehold-binding region and an 8-nt stem-invading region. Toehold-VISTA (Robson & Green, 2026; a machine-learning target-site model for conventional protein-output toehold switches) uses a 36-nt target site, comprising a 30-nt toehold-binding region and a 6-nt stem-invading region, and carries the RBS in the loop and the AUG in the stem. **b**, For each of the six held-out transcripts, measured $\log_2$(ON/OFF) of all tiled sites (grey) and of the 15 top- and 15 bottom-ranked sites predicted by DeSTAR (colored) and by Toehold-VISTA (grey); horizontal bars, group means; dashed line, $\log_2$(ON/OFF) = 0 (ON/OFF = 1). Across all six transcripts, DeSTAR's top-ranked sites were significantly more active than both the bottom-ranked sites and the transcript-wide average, whereas Toehold-VISTA's top-ranked sites were no more active than either (Mann-Whitney *U*-test, two-tailed; per-transcript statistics in Supplementary Table 4).

**a**

| Feature-class | Feature |
|---|---|
| Bottom 3-bp stem motif | Stem motif (S-S-W)<br>Stem motif (W-W-S)<br>Stem motif (S-S-S)<br>Stem motif (S-W-S)<br>Stem motif (W-S-S)<br>Stem motif (W-S-W)<br>Stem motif (W-W-W)<br>Stem motif (S-W-W) |
| Sensor folding ΔG | Hairpin ΔG<br>Toehold-hairpin ΔG |
| Target/context GC content | Target-window GC content<br>Seed-region GC content<br>Invasion-region GC content<br>Local-context GC content |
| Target accessibility | Mean single-base accessibility<br>Minimum single-base accessibility<br>Full target-window accessibility<br>Seed-region accessibility<br>Invasion-region accessibility |
| Relative transcript position | Relative transcript position |

**b**

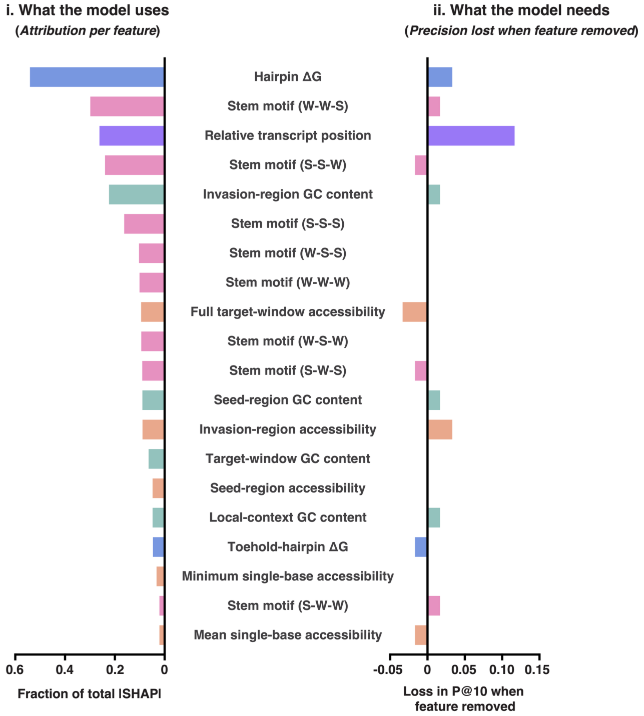


**c**

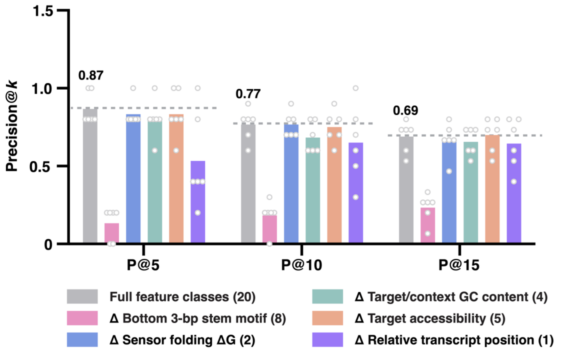


**d**

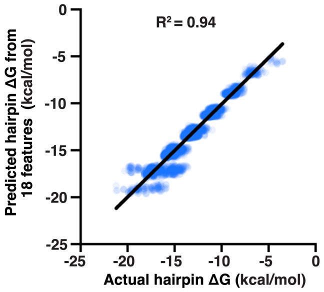


**e**

| Feature | Pearson's r with hairpin ΔG |
|---|---|
| Invasion-region GC content | -0.94 |
| Target-window GC conent | -0.49 |
| Stem motif (W-W-W) | 0.38 |
| Invasion-region accessibility | 0.37 |
| Stem motif (S-S-S) | -0.31 |
| Full target-window accessibility | 0.25 |

**Extended Data Fig. 5 | Feature attribution versus necessity in the DeSTAR model. a**, The 20 model features and their grouping into five feature classes: bottom 3-bp stem motif (eight one-hot strong/weak motifs), sensor folding ΔG (hairpin ΔG and toehold-hairpin ΔG), target/context GC content (four features), target accessibility (five features) and relative transcript position (one feature). **b**, Per-feature attribution and necessity. (i) *What the model uses*: mean absolute TreeSHAP value for each feature (ranking model trained on all six libraries, averaged over 10 seeds), expressed as a fraction of the total |SHAP| and ordered by attribution; hairpin ΔG is the single most-attributed feature, followed by the bottom-stem motifs. (ii) *What the model needs*: the loss in leave-one-transcript-out precision@10 when each feature is individually removed from the full model. Necessity and attribution diverge — most strikingly, hairpin ΔG has the highest attribution yet its removal barely changes precision, whereas relative transcript position causes a comparatively large loss. **c**, Leave-one-transcript-out precision@*k* (*k* = 5, 10, 15) for the full 20-feature model (grey; dashed line, full-model value) and for models with each feature class removed (Δ). Removing the bottom 3-bp stem motif class causes the largest drop at every *k*, confirming its necessity is consistent across precision cut-offs; full-model P@5/P@10/P@15 = 0.87/0.77/0.69. Open circles, per-transcript values; bars, mean; *n* = 6 transcripts. **d**, Actual hairpin ΔG versus hairpin ΔG predicted from the other 18 (non-ΔG) features (gradient-boosted regression trees, 5-fold cross-validation; $R^2 = 0.94$, $n = 7{,}383$ sensors). Hairpin ΔG is almost fully recoverable from the sequence-composition, GC and accessibility features — explaining its high attribution but negligible necessity in (**b**). **e**, Pearson correlation between individual features and hairpin ΔG ($n = 7{,}383$ sensors); invasion-region GC content is the strongest single correlate (r = -0.94).

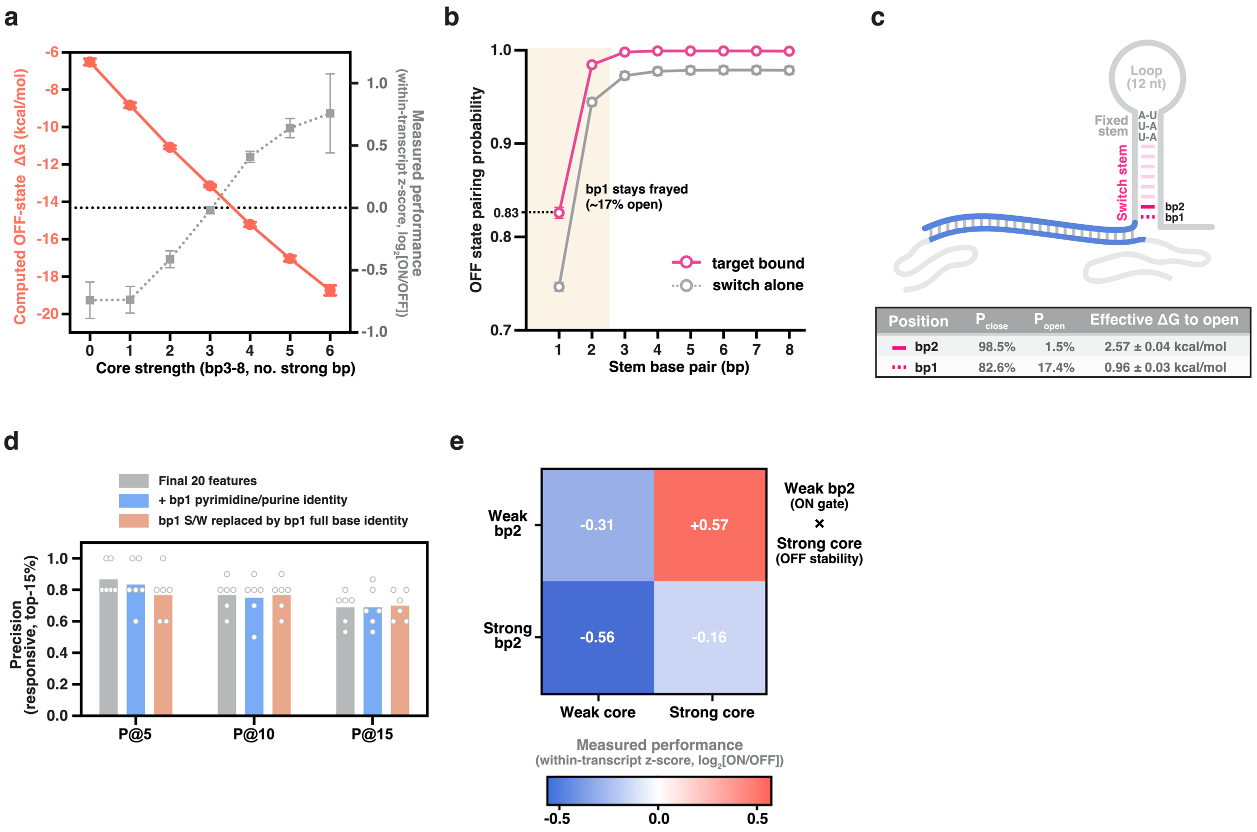


**Extended Data Fig. 6 | A biophysical rationale for the learned bottom-stem grammar. a**, Computed OFF-state hairpin free energy of the target-free switch (ViennaRNA 2.7, RNAfold minimum free energy, 37 ºC; left axis, orange) versus core strength (bp3-bp8, number of strong G·C pairs), overlaid with measured performance (within-transcript z-score of $\log_2$(ON/OFF); right axis, grey). Strengthening the core monotonically deepens the OFF-state hairpin, and measured ON/OFF rises in parallel. Points, mean; error bars, s.e.m.; $n$ = 6 transcripts. **b**, OFF-state base-pairing probability of stem pairs bp1-bp8 (ViennaRNA 2.7, partition function, 37 ºC) for the free switch (grey) and with the target bound to the toehold but stem invasion forbidden (magenta). The terminal pair bp1 is substantially frayed (open ~25% free, ~17% target-bound); pairing becomes reliable from bp2 onward. Points, mean; error bars, s.e.m.; $n$ = 6 transcripts. **c**, Top, schematic of the target-bound OFF state. The toehold is duplexed while the bp1-bp8 stem stays closed. Bottom, closed ($P_{close}$) and open ($P_{open}$) probabilities and the effective free energy to open, $\Delta G_{open} = RT \cdot \ln(P_{close}/P_{open})$ (RT = 0.616 kcal/mol at 37 °C), for bp1 and bp2 (mean ± s.e.m.; $n$ = 6 transcripts). $\Delta G_{open}$ is an effective, two-state, single-pair quantity; the s.e.m. reflects library-to-library reproducibility, not model uncertainty. **d**, Precision@$k$ at the responsive (top 15%) tier for three encodings of the toehold-terminal base pair (bp1) in the bottom-stem motif, for $k$ = 5, 10, and 15: the strong/weak encoding used in the model (Final 20 features); the same features plus a bp1 pyrimidine/purine feature (+ bp1 pyrimidine/purine identity); and bp1's strong/weak replaced by its full four-base identity within the motif (bp1 S/W replaced by full base identity). bp2, bp3 and all other features are identical across encodings; for each held-out transcript, candidate scores were averaged across the 10 seeds into a single consensus ranking before computing precision. Bars, mean across the six held-out transcripts; circles, individual transcripts. **e**, Mean within-transcript performance for sensors binned by bp2 identity (weak versus strong) and core strength (number of strong pairs among bp3-bp8: weak, 0-2; strong, 3-6). Cells, mean; $n$ = 6 transcripts.

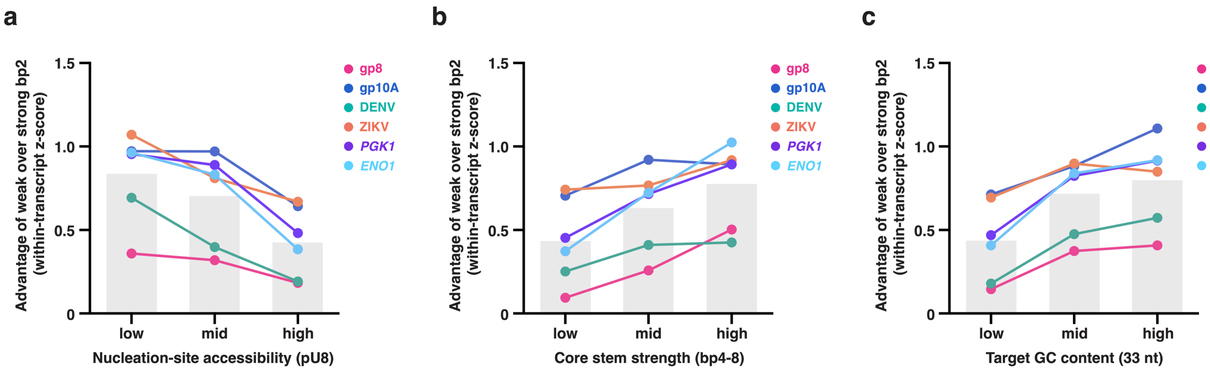


**Extended Data Fig. 7 | The weak-bp2 advantage is context-dependent.** In each panel, the weak-over-strong-bp2 advantage (mean within-transcript performance of weak-bp2 sensors minus that of strong-bp2 sensors, where performance is the within-transcript z-score of $\log_2$(ON/OFF)) is plotted against a biophysical covariate binned into low/mid/high terciles, for all six transcripts (colored lines) with the across-transcript mean (grey bars). **a**, Versus nucleation-site accessibility (pU8; RNAplfold unpaired probability of the 8-nt invasion segment). **b**, Versus core stem strength (number of strong G·C pairs in bp4-8; bp3 omitted to separate its tunning role from the core's OFF-state-stability contribution. **c**, Versus target GC content (across the 33-nt target site). Each dependence was consistent across all six transcripts; in a joint linear model of the weak-bp2 advantage on all three covariates (within-transcript z-scored, site level), each remained significant when adjusted for the other two (weak-bp2 × covariate interaction: accessibility $\beta$ = -0.08, $P = 6.0 \times 10^{-4}$; core $\beta$ = +0.09, $P = 2.5 \times 10^{-4}$; GC $\beta$ = +0.12, $P = 5.4 \times 10^{-7}$; $n$ = 7,369 sites, with 14 terminal sites lacking a defined pU8 excluded; Supplementary Table 4). Lines, individual transcripts; grey bars, across-transcript mean; $n$ = 6 transcripts.
